# Structural and genetic diversity in the secreted mucins, *MUC5AC* and *MUC5B*

**DOI:** 10.1101/2024.03.18.585560

**Authors:** Elizabeth G. Plender, Timofey Prodanov, PingHsun Hsieh, Evangelos Nizamis, William T. Harvey, Arvis Sulovari, Katherine M. Munson, Eli J. Kaufman, Wanda K. O’Neal, Paul N. Valdmanis, Tobias Marschall, Jesse D. Bloom, Evan E. Eichler

## Abstract

The secreted mucins MUC5AC and MUC5B play critical defensive roles in airway pathogen entrapment and mucociliary clearance by encoding large glycoproteins with variable number tandem repeats (VNTRs). These polymorphic and degenerate protein coding VNTRs make the loci difficult to investigate with short reads. We characterize the structural diversity of *MUC5AC* and *MUC5B* by long-read sequencing and assembly of 206 human and 20 nonhuman primate (NHP) haplotypes. We find that human *MUC5B* is largely invariant (5761-5762aa); however, seven haplotypes have expanded VNTRs (6291-7019aa). In contrast, 30 allelic variants of *MUC5AC* encode 16 distinct proteins (5249-6325aa) with cysteine-rich domain and VNTR copy number variation. We grouped *MUC5AC* alleles into three phylogenetic clades: H1 (46%, ∼5654aa), H2 (33%, ∼5742aa), and H3 (7%, ∼6325aa). The two most common human *MUC5AC* variants are smaller than NHP gene models, suggesting a reduction in protein length during recent human evolution. Linkage disequilibrium (LD) and Tajima’s D analyses reveal that East Asians carry exceptionally large *MUC5AC* LD blocks with an excess of rare variation (p<0.05). To validate this result, we used Locityper for genotyping *MUC5AC* haplogroups in 2,600 unrelated samples from the 1000 Genomes Project. We observed signatures of positive selection in H1 and H2 among East Asians and a depletion of the likely ancestral haplogroup (H3). In Africans and Europeans, H3 alleles show an excess of common variation and deviate from Hardy-Weinberg equilibrium, consistent with heterozygote advantage and balancing selection. This study provides a generalizable strategy to characterize complex protein coding VNTRs for improved disease associations.

## INTRODUCTION

Mucosal linings serve a dynamic role at the interface between internal tissues and the external environment. In the lumen of the lungs, epithelial cells provide defensive functionalities through mucociliary clearance, a mechanism in which mucus secretions trap inhaled pathogens for mechanical removal^1^. The secreted mucins *MUC5AC* and *MUC5B* are major components of mucus that contribute to its barrier function and act as receptor decoys for pathogens, such as the influenza virus that binds directly to mucin sialic acids^2^. These polymeric glycoproteins thus provide a critical innate immunological role in defending the airways against environmental insults; however, they have also been implicated in the pathogenicity of muco-obstructive airway diseases like asthma and cystic fibrosis^3^.

Despite their fundamental roles in maintaining epithelial homeostasis, *MUC5AC* and *MUC5B* sequence variation remains poorly understood. The challenge in assessing these loci is that they harbor large central exons (60-80% of total coding sequence) composed of variable number tandem repeats (VNTRs). These VNTRs encode numerous serine and threonine residues that are decorated with sialic acid, a terminal sugar moiety that is bound by the glycoproteins of some viral pathogens^2,4^. Limitations of short-read sequencing in assembling these repetitive loci have hindered efforts to accurately resolve copy number variation^5,6^. VNTR structural variants may affect the functional ability of mucins to act as barriers to pathogens and change their biochemical and biophysical properties; therefore, it is critical that the sequences of these loci in many human genomes are characterized to discover the most common patterns of variation directly affecting protein function for these important molecules.

Long-read sequencing technologies allow for the characterization of *MUC5AC* and *MUC5B* with haplotype-level resolution. Previously, gene references for both genes were constructed using Pacific Biosciences (PacBio) single-molecule, real-time (SMRT) sequencing from a limited number of humans. Four genome assemblies were used to characterize three distinct *MUC5AC* haplotypes for VNTR structural variation^7^. However, analyses of *MUC5AC* allele sizes via Southern blot suggest a much greater extent of human diversity^8^. Many additional human genomes have recently been sequenced with newer and more accurate high-fidelity (HiFi) circular consensus sequencing (CCS) as part of the Human Genome Structural Variation Consortium (HGSVC)^9^ and the Human Pangenome Reference Consortium (HPRC)^10^. Here, we leverage the large-scale sequencing efforts of the HGSVC and HPRC to explore common patterns of genetic variation in *MUC5AC* and *MUC5B,* specifically within the VNTR portion of the molecule. Using 206 diverse human haplotypes assembled with high-quality PacBio HiFi CCS reads, we characterize the genetic diversity of these loci in different human populations. We also compare the human alleles of *MUC5AC* and *MUC5B* to that of five nonhuman primate (NHP) species (chimpanzee, bonobo, gorilla, orangutan, and gibbon) to distinguish human-specific patterns of variation. Finally, we explore methods to genotype these loci using haplotype tagging single-nucleotide polymorphisms (tSNPs) and a structural variant genotyping tool. These results provide the first comprehensive view of VNTR variation and evolution in the secreted airway mucins *MUC5AC* and *MUC5B* and outline a path forward for improved disease association studies.

## METHODS

### Long-read sequence assembly and QC

Whole-genome assemblies from 104 HPRC^10^ (n = 47) and HGSVC ^9^ (n = 57) samples were leveraged for *MUC5AC* and *MUC5B* variant discovery. These genomes include 49 Africans, 23 Admixed Americans, 14 East Asians, 10 Europeans, and 8 South Asians (Table S1). Sequencing for both cohorts was conducted using PacBio HiFi CCS. Average HPRC sequencing coverage was 42× (minimum = 31×) and average HPRC read N50 was 19.7 kbp (minimum = 13.5 kbp). Average HGSVC sequencing coverage was comparable at 40× (minimum = 25×) and average read N50 was 17.2 kbp (minimum = 10.0 kbp). The HPRC genome assembly was conducted by Liao et al.^10^ using Trio-Hifiasm^11^ (maternal and paternal short reads used in haplotype phasing). We assembled 54 HGSVC samples using Hifiasm v0.16.1^11^ (pseudo-haplotype resolved phasing). For the remaining three HGSVC samples with trio information (HG00514, HG03125, NA12878), we used paternal and maternal short reads with yak v0.1 (https://github.com/lh3/yak) to create k-mer databases for contig phasing in the child’s assembly with Hifiasm v0.15.1^11^ (see Ebert et al.^9^ for parental short-read information). Average HPRC haplotype assembly N50 was 40.8 Mbp (minimum = 17.4 Mbp) and average HGSVC haplotype assembly N50 was 55.2 Mbp (minimum = 14.1 Mbp). Regional assembly contiguity and reliability for the *MUC5AC/5B* locus was assessed using the flagger pipeline^9^ and Nucfreq, a method to detect potential misassemblies and collapses in phased haplotypes^12^. We also inspected for assembly misalignments in the locus using SafFire (https://github.com/mrvollger/SafFire).

We assessed 10 total NHP genome assemblies for chimpanzee (n = 2), bonobo (n = 2), gorilla (n = 2), Sumatran orangutan (n = 2), Bornean orangutan (n = 1), and Siamang gibbon (n = 1). Specifically, these included PTR1 (Central chimpanzee, Clint), PPA1 (bonobo, Mhudiblu), GGO1 (Western gorilla, Kamilah), and PAB1 (Sumatran orangutan, Susie) haplotype-resolved assemblies for *MUC5AC/5B* assembled with Hifiasm v0.15.1^13^. All other NHP assemblies were generated as part of the primate T2T (telomere-to-telomere) Consortium and assemblies were downloaded from GenomeArk^14^; these include PTR2 (Central chimpanzee, AG18354), PPA2 (bonobo, PR00251), GGO2 (Western gorilla, Jim), PAB2 (Sumatran orangutan, AG06213), PPY1 (Bornean orangutan, AG05252), and SSY (Siamang gibbon, Jambi). These assemblies were constructed using both high-coverage PacBio HiFi CCS reads and ultra-long (UL) Oxford Nanopore Technologies (ONT) reads via the Verkko 2.0^15^ assembler. Information about assembly quality and validation can be found in Mao et al.^13^ and Makova et al.^14^ We inspected the *MUC5AC/5B* regional assembly contiguity using SafFire in the same manner as the human HGSVC assemblies.

### Sequence extractions and phylogenetic analyses

HPRC, HGSVC, and NHP phased genome assemblies were aligned to CHM13^16^ using minimap2 v2.24^17^ with CIGAR string inclusion, full-genome alignment divergence less than 10%, secondary alignments suppressed, and a minimal peak all versusf all alignment score of 25000. Coordinates for a specific locus in individual haplotype assemblies were identified using rustybam v0.1.29 (https://github.com/mrvollger/rustybam) and sequences were extracted using seqtk v1.3 (https://github.com/lh3/seqtk). Exon and intron boundaries were defined based on human GENCODE V35^18^ annotations in CHM13^16^ (MUC5AC-201, MUC5B-204). Intronic and flanking intergenic sequences used to construct phylogenies were selected in a recombination-aware manner, based on UCSC Genome Browser 1000 Genomes Project (1KG) linkage disequilibrium (LD) structure annotations^19^. A multiple sequence alignment (MSA) was conducted using MAFFT v7.487^20^ with global pairwise alignment and 100 iterations, followed by visual inspection of alignment quality using Jalview v9.0.5^21^. Segments of the MSA determined to be misaligned were identified and eliminated manually. Maximum-likelihood tree calculations were performed using iqtree v1.6.12^22^ with automatic model selection and 1,000 bootstraps. All phylogenetic trees in figures were constructed using ggtree v3.2.1^23^ in R v1.4.2 (https://www.R-project.org). Haplogroup coalescence times were estimated with iqtree2^24^ based on estimated chimpanzee divergence (6.4 million years ago [mya])^25^.

### Gene and protein domain/VNTR motif annotations

Computational protein prediction for all human and NHP haplotypes was conducted via the same alignment pipeline as phylogeny construction based on human exon annotations from CHM13^16^. We predicted translated exons using the ExPasy tool in EMBOSS v6.6.0^26^. For computational protein predictions that were complete (i.e., complete open reading frame [ORF], no truncations), protein domain annotations were manually curated using cys domain and VNTR domain sequences annotated previously by Guo et al.^7^ Protein groups (P1-P6) were defined for *MUC5AC* as containing more than one haplotype and variation in cys domain copy number, tandem repeat domain copy number, and/or repeat motif copy number variation in homologous VNTR domains. Protein groups for *MUC5B* were similarly defined; however, the inclusion criteria of harboring more than one haplotype per group was dismissed due to protein sequence length variation in three singletons for *MUC5B* (P1, P4, P5). We characterized motif variation across individual VNTR domains for human *MUC5AC* and *MUC5B* based on previously published consensus motif sizes (24bp/8aa for *MUC5AC*^27^, 87bp/29aa for *MUC5B*^28^). Heatmaps of motif usage for all haplotypes of *MUC5AC* and *MUC5B* were constructed using a custom R script that included normalization on total VNTR sequence space (motif counts / total number of motifs) to account for length variability, normalization within motifs, and hierarchical clustering (Unweighted Pair Group Method of Arithmetic Mean [UPGMA] clustering^29^) of haplotypes and motifs for group visualization. Similarly, motif diagrams in linear sequence space were constructed using a custom R script that designated a unique color to each distinct motif and clustered unique alleles by row using UPGMA.

### NHP allele alignments and intronic VNTR analysis

We generated all versus all alignments between the most common haplotypes of *MUC5AC* and *MUC5B* in humans and NHPs using minimap2^17^ with the same parameters as phylogenetic analyses. Tiled alignment plots for each locus were constructed using SvbyEye v0.99.0 (https://github.com/daewoooo/SVbyEye) in R v4.3.1 with a bin size of 10,000 bp and custom percent identity breaks. VNTR sequences in intron 15 and ∼3kb before the start codon of *MUC5AC* were curated using tandem repeats finder v4.10^30^ with the following parameters: match = 2, mismatch = 7, delta = 7, PM = 80, PI = 10, minimum alignment score = 50, and max period size = 30. Detection of H3 k-mers for the intronic VNTR was conducted using STREME from the MEME suite of motif-based sequence analysis tools v5.5.4^31^.

### LD block structure and selection detection analyses

Illumina whole-genome sequencing (WGS) data from the most recent high-coverage (30×) release of the 1KG^19^ were used to assess the LD structure of the *MUC5AC/MUC5B* locus. These data include WGS from 2,600 unrelated individuals: 691 African, 526 European, 514 South Asian, 515 East Asian, and 354 American genomes. LDBlockShow v1.40^32^ was used to construct LD plots based on D′ ^33^ for all single-nucleotide polymorphisms (SNPs) in the *MUC5AC/MUC5B* region (GRCh38 coordinates, chr11:1117952-1272172). Autosome-wide LD block calculations were estimated with the PLINK v1.9^34^ blocks parameter, which estimates haplotype blocks based on definitions described by Gabriel et al^35^ (the region of chromosome 11 that harbors *MUC5AC* and *MUC5B* features a high recombination rate)^36^. Calculations were limited to SNPs with a minor allele frequency greater than 5%, those with 75% or higher genotyping rate, and those in Hardy-Weinberg equilibrium. To assess if the region of chromosome 11 containing *MUC5AC* and *MUC5B* showed signatures of selection, Tajima’s D^37^ analysis was conducted using the phased 1KG cohort of samples. Relative to an autosome-wide distribution, significantly positive values of Tajima’s indicate there is an excess of high-frequency variation (suggestive of balancing selection), while significantly negative values indicate there is an excess of rare variation (suggestive of positive selection). Calculations were computed for all autosomes and were specific to the five global populations. Each chromosome was partitioned into 10 kbp bins with filtering for bins that contained at least 10 SNPs. Tajima’s D statistics were computed for bins using PLINK v1.9^34^ and regions harboring significant signatures of either positive or balancing selection were based on the 90^th^ and 95^th^ percentiles of values in the autosome-wide distribution (significantly negative Tajima’s D is suggestive of positive selection, significantly positive Tajima’s D is suggestive of balancing selection).

### New tagging SNPs (tSNPs) and mapping of disease relevant GWAS SNPs

To uncover SNPs in significant LD with VNTR haplogroups of *MUC5AC* and *MUC5B,* phylogenetic haplogroups from the HGSVC/HPRC genomes were encoded as biallelic SNPs. Calculation of squared correlations between these variants encoding haplogroup identity and all SNPs within 50 kbp of the loci were performed using PLINK v1.9^34^. Genome-wide association study (GWAS) risk alleles for *MUC5AC* and the phenotypes of asthma/allergy and infection-induced pneumonia/meningitis were mined through the GWAS catalog. Variants were included in subsequent LD analysis if they had a reported p-value of 1x10^-9^ or smaller for the phenotype association, had the nucleotide annotation for the risk allele, and were unambiguously mapped to the HPRC/HGSVC genomes. The final set of variants included six SNPs from six GWAS studies (rs35225972^38^, rs11245962^39^, rs28415845^40^, rs11245979^41^, rs28737416^42^, and rs28729516^43^). Squared correlation values were calculated in the same manner as tSNP discovery.

### Genotyping of *MUC5AC* haplogroups in 1KG populations using Locityper

*MUC5AC/5B* genotyping was performed with Locityper v0.10.9 (https://github.com/tprodanov/locityper) and its dependencies SAMtools v1.19^44^, jellyfish v2.3.0^45^, and strobealign v0.11.0^46^. Diploid genomes from the HGSVC/HPRC sample set were included as alleles in the reference panel if they were complete for the *MUC5AC/5B* locus (no assembly breaks or alignment ambiguities), annotated for both haplogroups, and had accessible high-quality short reads through the 1KG dataset. The final set of genomes that constituted the reference panel included 99 genomes (i.e., 198 haplotypes) for both *MUC5AC* and *MUC5B* that were sequence curated for haplogroups/protein groups and had high-quality paired-end short-read data.

CHM13^16^ was used as the reference genome for all Locityper analyses, with gene coordinates set to chr11:1227366-1274380 and chr11:1292367-1334784 for *MUC5AC* and *5B*, respectively. For leave-one-out (LoO) analyses, the target sample for genotyping was excluded from database construction and the highest alignment accuracy level was used. All other options for database construction, sequencing dataset preprocessing, and genotyping were set to default. Genotyping accuracy was determined based on edit distance (alignment differences) between the real and retrieved genotypes during LoO and compared to the closest “available” genotype (smallest edit distance between true genotype and all possible diploid combinations of alleles in the reference panel). Computation of edit distances between alleles in the LoO concordance analysis was performed using the Locityper helper script “gt_dist.py.”

### *MUC5AC* and *MUC5B* Phenome-Wide Association Studies (PheWAS) in *All of Us*

Data from the *All of Us* Research Program^47^ controlled tier database were analyzed for a phenome-wide association study (PheWAS) with the *MUC5B* promoter polymorphism rs35705950^48^ and tSNPs for the major haplogroups of *MUC5AC* variants. As of January 2024, this cohort included ∼245,400 individuals with short-read WGS data, of which ∼185,000 were unrelated, annotated for age/sex, and had paired electronic health record (EHR) data (reported as International Classification of Diseases [ICD] codes). These individuals were categorized previously by the consortium for genetic ancestry using principal component analysis. We surveyed samples from African (AFR), European (EUR), East Asian (EAS), Admixed American (AMR), and Middle Eastern (MID) ancestries for *MUC5B* rs35705950 and tSNPs in high LD with *MUC5AC* haplogroups H1 (rs2075842, rs1132433, rs1132434, rs28652890, rs879136008), H2 (rs1015856541, rs28519516, rs28558973, rs28368633), and H3 (rs36154966, rs1004828576, rs940158763, rs36151150, rs36132281, rs35779873). We only included samples with genome quality scores ≥20 at individual loci; therefore, the final sample sets included ∼32,500 Africans, ∼3,200 East Asians, ∼2,000 South Asians, ∼98,600 Europeans, ∼28,200 Admixed Americans, and ∼650 Middle Eastern (MID), totaling ∼165,150 individuals (exact number of individuals varied between locus associations in respective populations; Tables S5 and S6). We included both ICD-9 and ICD-10 phenotype codes from patient EHRs and samples with male/female self-reported biological sex aged 20 years or older.

PheWAS analysis was performed using the R package PheWAS as outlined in Bick et al.^47^ The package translated ICD-10 codes to ICD-9 and calculated case and control genotype distributions, allelic p-value, and allelic odds ratio (OR) for each condition. A minimum count of two related codes was used to determine if a phenotype was sufficiently represented in the health data for association. Sex at birth, age at sample collection, and principal component analyses 1-3 were used as covariates. The aggregate.fun function was used to correct for duplicates in the EHR. Nominal p was set to < 2.7E-5 (p-adjusted < 0.05 after Bonferroni correction) for phenotype associations with rs35705950 and *MUC5AC* tSNP alleles in the dataset.

## RESULTS

### *MUC5AC/5B* assembly and QC

We targeted assembly and quality control (QC) of a ∼160 kbp region including *MUC5AC* and *MUC5B* from 104 human genomes. This included 47 genomes from the HPRC and 57 from the HGSVC where long-read sequencing data had recently been generated and made publicly available^9,10^. To harmonize genome assembly, we generated phased genome assemblies using the same computational pipeline used for the generation of HPRC assemblies (Methods) from HiFi PacBio sequencing data. The combined sample set includes 49 Africans, 23 Admixed Americans, 14 East Asians, 10 Europeans, and 8 South Asians (Methods & Table S1). Next, we applied the flagger^10^ and Nucfreq^12^ computational pipelines to detect collapses or misassemblies across the 160 kbp target region. Of the 208 total human haplotypes, 206 (99%) were classified as correctly assembled without gaps, breaks, or misjoins in the *MUC5AC/5B* region. Two haplotypes (one each) from samples HG01114 and HG02509 were fragmented and excluded from all downstream analyses. For comparative purposes, we performed a similar analysis from 11 individuals from six NHP species for which HiFi sequencing data have recently been generated^13,14^ (Table S1). All *MUC5AC* loci passed QC with no ambiguous alignments to CHM13; in contrast, three NHP assemblies for *MUC5B* failed to pass QC, namely, one gorilla haplotype (Kamila h2) and both haplotypes of a Sumatran orangutan (Susie h1 and h2). Therefore, these NHP haplotypes were removed from further *MUC5B* analyses (see below).

### Human MUC5AC protein and genetic diversity

To understand human genetic diversity in *MUC5AC*, we first constructed a phylogeny centered around the gene model. We extracted 26.5 kbp of noncoding sequence flanking *MUC5AC* exons for the 206 human haplotypes and generated a maximum likelihood phylogenetic tree using chimpanzee as an outgroup. Human alleles were grouped into three distinct haplogroups or clades (Fig. 1a), namely H1 (n=103), H2 (n=78), and H3 (n=25). H1 is the most phylogenetically distinct (100 bootstrap support), is reduced in frequency among African genomes (p=4x10^-3^ comparing H1 to H2/H3 frequencies via chi-square, Fig. 1b), and is estimated to have arisen most recently. For example, we estimate an H1 coalescent of ∼120,000 years ago when compared to H2 or H3 (∼330,000 years ago).

Next, we predicted the protein gene model associated with each human haplotype (Methods). We identified 16 distinct MUC5AC protein variants with extensive length variation (Fig. 1c). The three most common protein variants, 5654 aa/96 haplotypes, 5742 aa/67 haplotypes, and 6325 aa/15 haplotypes (Fig. 1c) project onto the phylogenetic haplogroup designations H1, H2, and H3, respectively. There is, however, additional variation not immediately apparent from the phylogeny that instead is discovered through detailed protein curation of sequence from the large central exon. Guo et al.^7^, for example, classified protein variants into three groups (P2, P3 and P6) based on MUC5AC domain annotations. We extend this classification by identifying three additional protein variant groups (P1, P4 and P5) based on VNTR domain, cys domain, and VNTR motif copy number variation.

**Figure 1.**
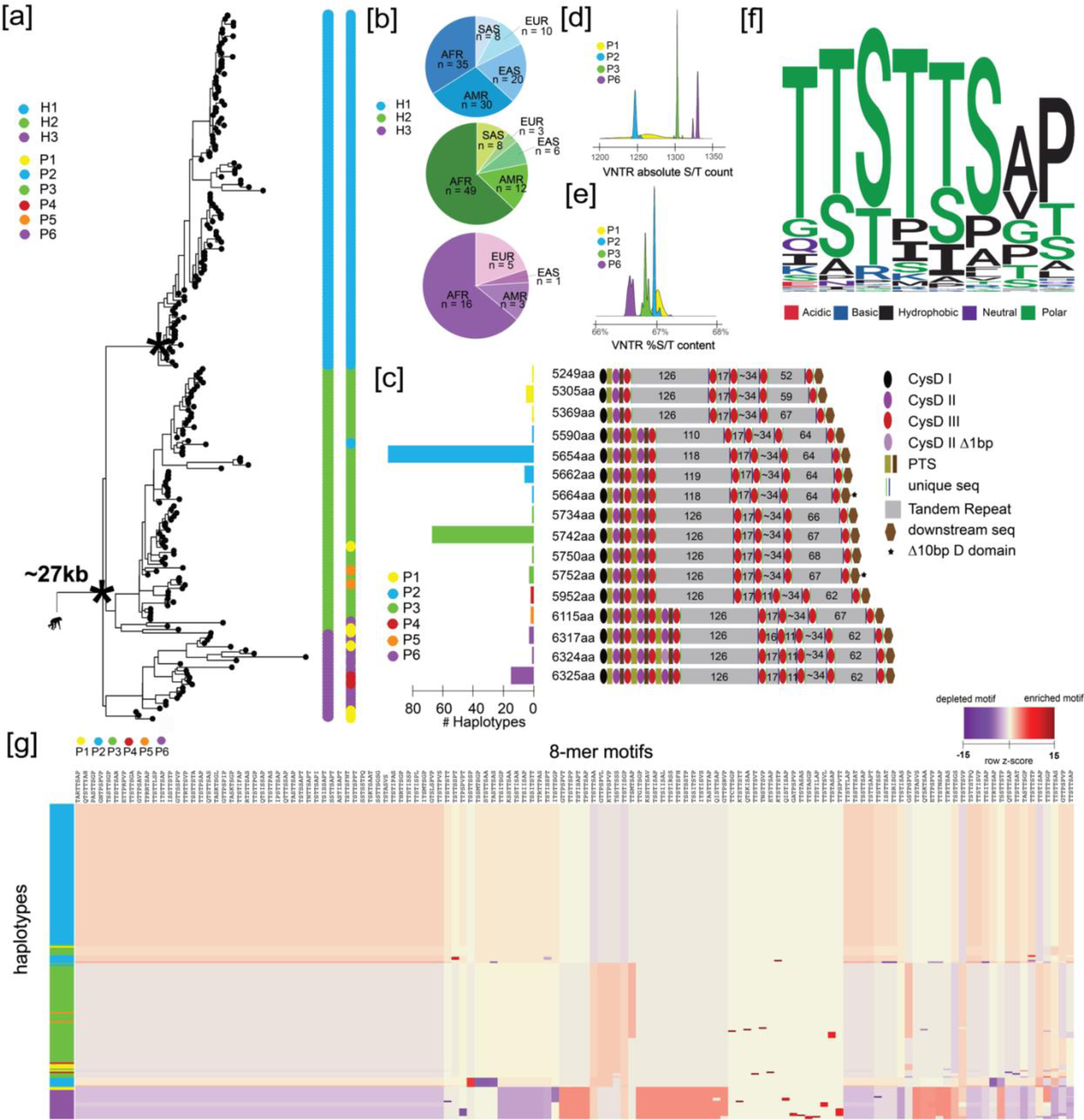
The genetic architecture of *MUC5AC* in 206 human haplotypes. **(a)** Recombination-aware phylogenetic analysis of ∼25 kbp neutral sequence (5.592 kbp from introns 31-48 and 21 kbp from 3’ flanking sequence) from 206 human haplotypes of *MUC5AC* with two chimpanzee haplotypes as outgroup. (*) = central node with 100 bootstrap support. H1-H3 correspond to three major haplogroups; P1-P6 correspond to protein groups (consistent with panel c). **(b)** Frequency of population-specific haplotypes found in the three major phylogenetic haplogroups of *MUC5AC.* AFR = African, AMR = American, EAS = East Asian, EUR = European, SAS = South Asian. **(c)** Protein predictions for haplotypes of *MUC5AC.* Diagrams represent protein domains with the large central exon of *MUC5AC* and modeled after Guo et al.^7^ Text colors correspond to protein groups visualized in panel a. **(d)** Absolute serine and threonine (S/T) count across variable number tandem repeat (VNTR) domains within the four most common protein groups of MUC5AC. **(e)** Percent S/T content within VNTR domains for the four most common protein groups of MUC5AC. **(f)** Logo plot of the 130 8-mer amino acid motif variants used in MUC5AC VNTR domains. **(g)** Heatmap of 8-mer motif utilization across 206 protein variants of human *MUC5AC.* Heatmap constructed with normalization within motifs (columns) and hierarchical clustering of haplotypes (rows) and motifs (columns).

Considering the domain architecture, most MUC5AC protein variants harbor four distinct tandem repeat domains (P1-3, P5); however, two groups (P4, P6) harbor an additional central tandem repeat domain with 11 copies of the MUC5AC 8-mer repeat motif and an additional cys domain. Most MUC5AC protein variants also feature five cys domains preceding the first tandem repeat domain; however, variants in P5/6 harbor a duplication containing type 2 and type 3 domains, while variants in P1 harbor a novel deletion of these domains. Motif copy number within individual VNTR domains is also extensive in the first and last domains in each protein variant group.

We characterized the composition of the MUC5AC degenerate VNTR 8-mer repeat because the density of serines and threonines is critical for mucin barrier function and provides numerous sites for potential glycosylation and phosphorylation. We find that the absolute count of serine and threonine residues across the VNTR domains correlates positively with protein length (Fig. 1d); however, when normalized for the total length of the VNTR, the two shortest protein variant groups (P1 and P2) harbor the highest concentration of serines and threonines (Fig.1e). There are a remarkable 211 unique 24-mers (nucleotides) and 130 unique protein 8-mer motifs (amino acids) diversifying the protein coding region across 206 human haplotypes. Motif changes, however, are constrained, with most harboring the pattern of TTSTTS in the first six amino acids (Fig. 1f, Fig. S1a). The preferential use of threonines in many of these motifs is likely a consequence of the higher propensity for threonines to harbor O-glycans relative to serines^49^, facilitating extensive binding potential of viral glycoproteins to MUC5AC. Furthermore, the high incidence of prolines in these motifs likely contributes to the glycosylation potential of nearby serines/threonines by exposing these residues in a β-turn conformation^50^.

Of the 130 unique protein 8-mer motifs for MUC5AC, only nine are unique to a single haplotype, indicating that much of this variation is often shared between protein isoforms. There are distinctive modules of motifs that cluster together in frequency of usage for protein groups 2, 3, and 6 (Fig. 1g). Those harboring the most distinctive usage signature include variants in P6, while those in P2 are consistently homogenous. When considering all tandem repeat domains, we identify 30 unique alleles of *MUC5AC* that cluster predominantly within their phylogenetic haplogroups (Fig. S1a). Most motif variation is due to nonsynonymous amino acid changes between haplotypes; however, there are instances where entire motifs have been gained or lost. Such structural changes explain the absence of 7-10 motifs in TR1 and 3 motifs in TR5 for H1 haplotypes, as well as the absence of 4 motifs in TR5 common to H3 haplotypes. Overall, there is extensive cys domain copy number, VNTR copy number, and VNTR motif usage variation in the large central exon of *MUC5AC* across human haplotypes.

### Human *MUC5B* genetic and protein diversity

Similarly, we repeated the analysis for the *MUC5B* locus and observed far less genetic and protein variability for this locus when compared to *MUC5AC*. A maximum likelihood phylogenetic tree (24.6 kbp intronic sequence using chimpanzee as an outgroup) distinguishes two distinct human haplotypes with 100 bootstrap support (Fig. 2a). The most common haplogroup, H2, was identified in 82% (169/206) of assembled haplotypes and is estimated to have emerged ∼770,000 years ago, while the less abundant H1 (18%) predictably arose more recently (∼407,000 years ago). While H2 is found across all continental populations, H1 notably shows a reduced frequency in populations of East Asian descent (Fig. 2b). At the protein level, we predict a complete ORF for 92% (190/206) of the assemblies while 16 predict a premature stop codon (Fig. 2c). Owing to the large number of homopolymers associated with the VNTR in the large central exon of MUC5B (exon 31), we hypothesized that these 16 haplotypes with disruptions in their ORF arose because of assembly artifacts. To test this, we reassembled eight of these samples where both ONT and HiFi sequence data were available and applied a different assembly algorithm (Verkko^15^). Reassembly recovered and re-established the ORF for three with predicted protein lengths consistent with those predicted for the representative haplotypes.

Among the 190 haplotypes with complete ORFs, 87% predict proteins with the canonical MUC5B length of 5762 aa (P3). The second most abundant, P2, differs in length by one amino acid (5761 aa) and represents 9% of protein isoforms. This slight change associates exclusively with the H1 haplogroup. These findings support the long-standing belief that *MUC5B* is less variable than *MUC5AC*^51^. Our deeper survey, however, suggests that the locus is not invariant. We identify seven haplotypes (3.6%, 7/190 complete proteins) where the protein is predicted to have elongated (6291-7019aa, P4-P6) due to expansion of the VNTR domains. Five of these longer variants harbor seven total VNTR domains with an excess of ∼800 amino acids of tandem repeat sequence and two additional cys domains. Unlike *MUC5AC,* there is no variation in cys domain copy number preceding the first tandem repeat domain in *MUC5B* variants. All seven elongated variants were found exclusively in individuals of African descent; therefore, much like *MUC5AC,* the ancestral state of this locus may have been longer.

**Figure 2.**
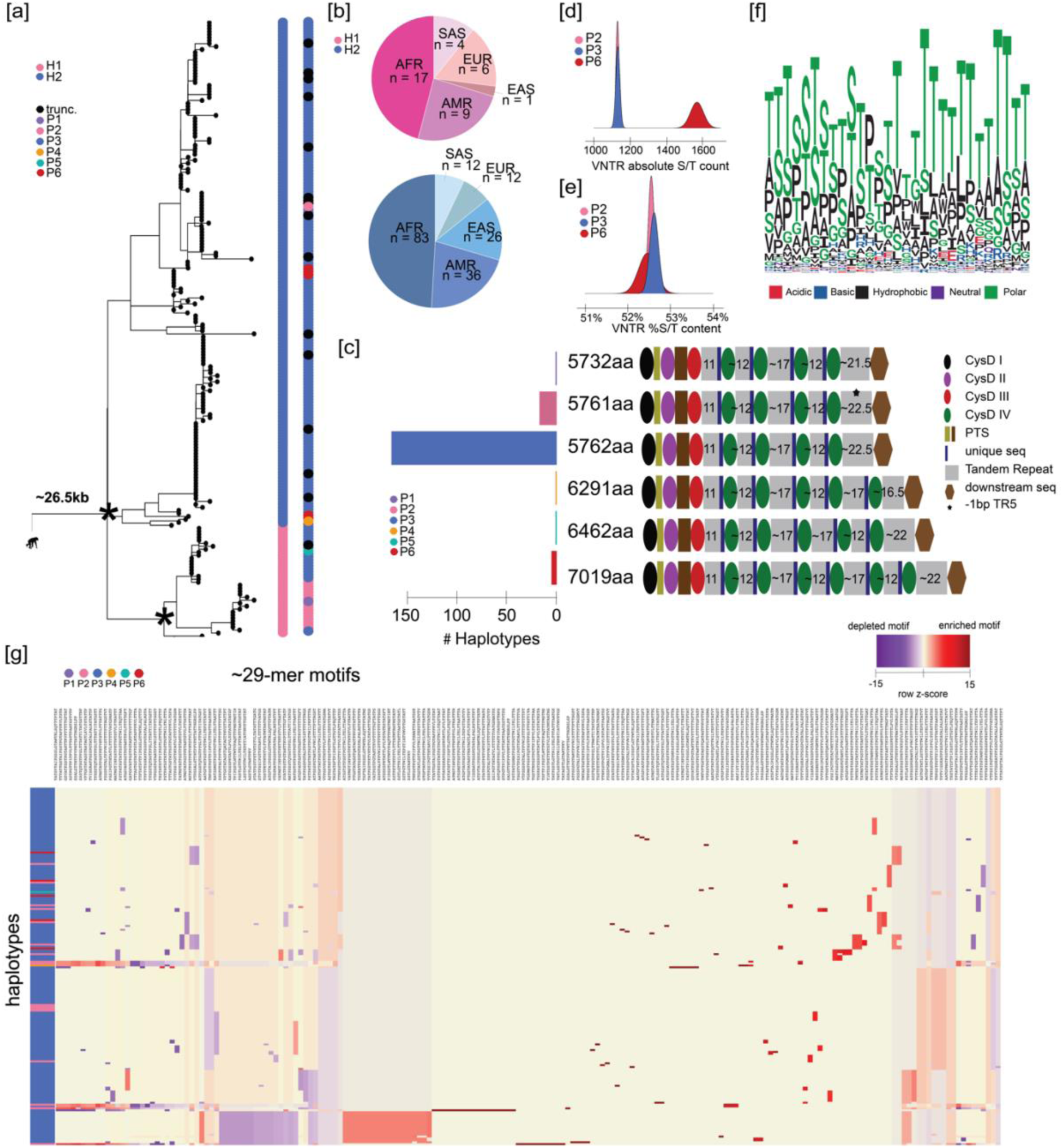
The genetic architecture of *MUC5B* in 206 human haplotypes. **(a)** Recombination-aware phylogenetic analysis of ∼26.5 kbp neutral sequence (introns 16-48) from 206 human haplotypes of *MUC5B* with two chimpanzee haplotypes as outgroup. (*) = central node with 100 bootstrap support. H1 and H2 correspond to two major haplogroups; P1-P6 correspond to protein groups (consistent with panel b); trunc. corresponds to haplotypes with truncated protein predictions. **(b)** Frequency of population-specific haplotypes found in the two major phylogenetic haplogroups of *MUC5B.* AFR = African, AMR = American, EAS = East Asian, EUR = European, SAS = South Asian. **(c)** Protein predictions for 206 human haplotypes of *MUC5B.* Diagrams represent protein domains with the large central exon of *MUC5B* and modeled after those in Ridley et al.^47^ Text colors correspond to protein groups visualized in panel a. **(d)** Absolute serine and threonine content (S/T) across VNTR domains for the three most common protein groups of MUC5B. **(f)** Logo plot of the complete 29-mer amino acid motif variants used in MUC5B VNTR domains across 206 human haplotypes. **(g)** Heatmap of 190 ∼29-mer motif utilization across protein variants of human MUC5B. Heatmap constructed through normalization for total VNTR sequence length, normalization within each motif (columns), and hierarchical clustering of haplotypes (rows) and motifs (columns).

The novel domains TR5 and TR6 associated with P4-P6 are most like TR3 and TR4, respectively, in repeat copy number and motif composition (Fig. S1b). P5 has compositionally unique versions of TR4 and TR5 that are more similar with motif usage to canonical TR4 and TR3, respectively. These results suggest that the acquisition of new tandem repeat domains has been accomplished via duplication of the central domains in *MUC5B*, rather than from the first and last domains. While the largest MUC5B protein isoform (P6) has increased in size due to VNTR expansion, it is interesting that serine and threonine abundance is relatively comparable to that of the more abundant canonical forms (P1-P4) (Fig. 2d-e). Like MUC5AC, threonine is favored across the irregular MUC5B repeat motif (Fig. 2f). Even though there are fewer distinct MUC5B protein variants, there are 191 unique 29-mers used across the haplotypes and 63 unique alleles across tandem repeat domains connected in linear sequence space (Fig. 2g, Fig. S1b). Unlike *MUC5AC,* there appear to be no gain or loss of whole motifs (i.e., motif variation is restricted to nonsynonymous mutations). Additionally, while P3 features unique modules of motif usage relative to other variants (Fig. 2g), frequency of motif usage is largely conserved across the different haplotypes of *MUC5B*.

### NHP variation in *MUC5AC* and *MUC5B*

We reconstructed the evolutionary history of *MUC5AC* and *MUC5B* by identifying orthologous loci from recently sequenced and assembled NHP genomes^13,14^. This included chimpanzee (n=2), bonobo (n=2), gorilla (n = 2), orangutan (n=3, Sumatran and Bornean species), and Siamang gibbon used as an outgroup (Fig. 3-4 and Table S1). For the *MUC5AC* locus, all NHP haplotypes (n = 22) predicted a complete ORF similar in structure to that of the human haplotypes and harbored both cys domain and VNTR variation (Fig 3a). Variation in the number of cys domains preceding the first tandem repeat domain is seen in alleles for chimpanzee and bonobo. One haplotype each from bonobo and gorilla harbor a truncated type I cys domain at the 5’ end of the exon that is not observed in any human, chimpanzee, orangutan, or gibbon haplotypes. Extensive variation in motif copy number and tandem repeat domain number was observed, with the Asian apes, orangutan, and gibbon carrying the longest predicted proteins. In fact, the most common protein variant in orangutan is approximately 1,500 amino acids longer than the longest human variant. All NHP variants were longer than the two most common human variants (H1 and H2), ranging in size from 6243aa-7887aa, due to increased exon 31 VNTR length (Fig. 3b). This suggests there has been a reduction of VNTR length in the human lineage (Fig. 3b-c).

Additionally, we characterized two noncoding VNTRs associated with the *MUC5AC* locus—an 8-mer VNTR mapping to intron 15 of *MUC5AC* that is unique in copy number and motif usage in human haplotypes (Fig S2, Fig. 3c and Note S1) and an 8-mer VNTR approximately 1-3 kbp in size, mapping upstream of the *MUC5AC* start codon. Based on ENCODE H3K27 mapping data^18^, the region corresponds to a potential enhancer. Diminished copy number variants of the enhancer VNTR have been associated with decreased *MUC5AC* expression^53^ and susceptibility to severe gastric cancer^54^. We find complete enrichment of shorter variants (less than 1,500 bp in length) in East Asians H1 haplotypes (Fig. S3) and an excess of long variants (greater than 2,000 bp) among African haplotypes in the HPRC/HGSVC sample set (X^2^ = 87.4, p<0.001). This suggests a potential founder effect or selection among East Asians that could result in population-specific differential expression of H1 *MUC5AC* variants. Additionally, all NHP haplotypes feature lengths of 881 bp-1649 bp (shortest in orangutan and longest in chimpanzee) for this enhancer VNTR.

**Figure 3.**
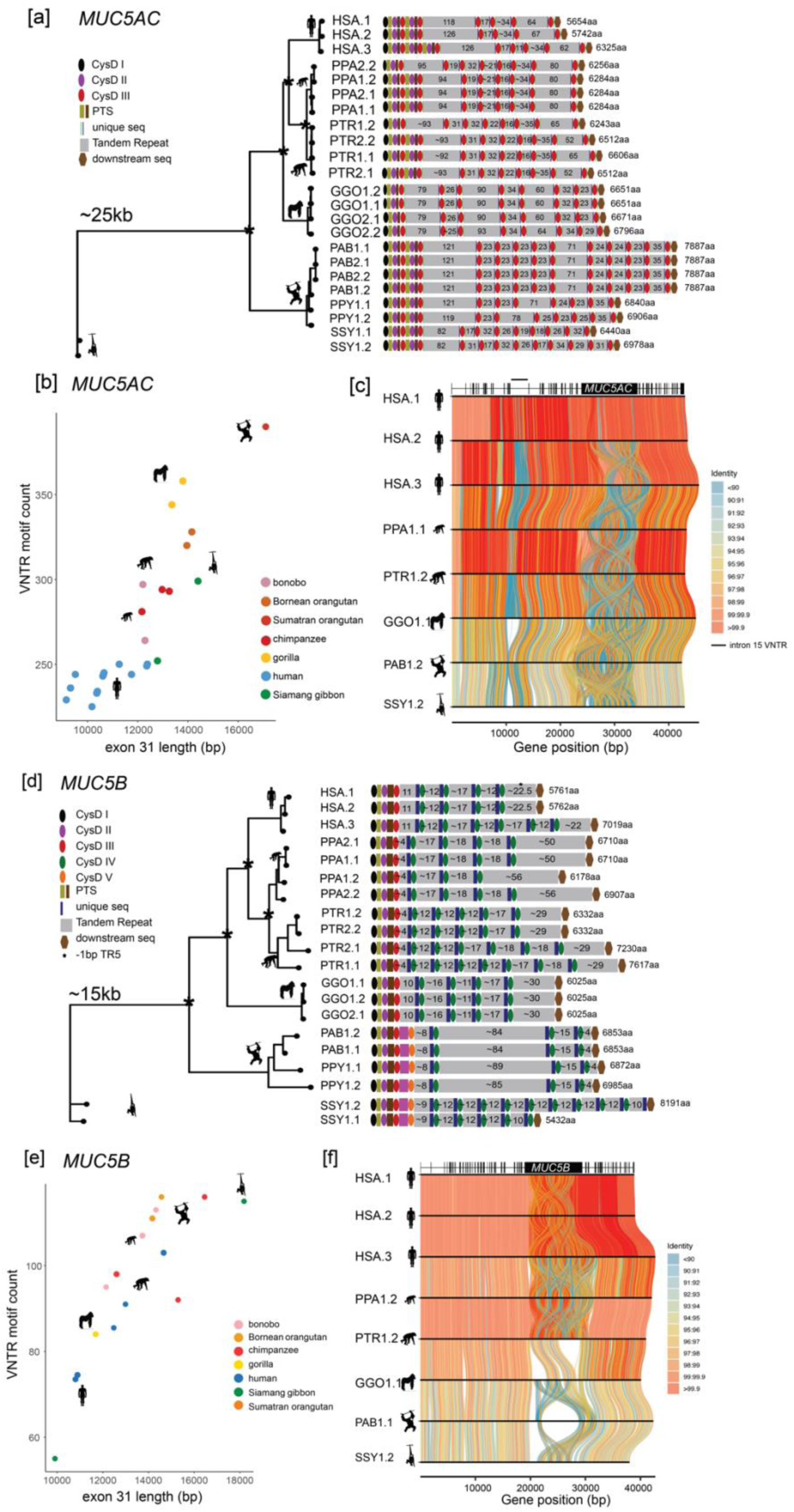
The genetic architecture of *MUC5AC* and *MUC5B* in the nonhuman ape lineages. **(a)** Phylogenetic analysis of ∼25 kbp from at minimum two haplotypes per ape lineage and subsequent protein predictions for *MUC5AC* haplotypes based on human exon boundary alignments. (*) = central node distinguishing species branches with 100 bootstrap support. Diagrams represent protein domains within the large central exon. HSA = human; PPA = bonobo; PTR = chimpanzee; GGO = gorilla; PAB = Sumatran orangutan; PPY = Bornean orangutan; SSY = Siamang gibbon. **(b)** Scatterplot of total *MUC5AC* exon 31 length (in base pairs) and total VNTR motif count across all VNTR domains in human and nonhuman primates. **(c)** Tiled alignments between representative haplotypes of each ape species (most common or most structurally unique haplotype per species) for *MUC5AC. MUC5AC* intron/exon boundaries are distinguished by the gene model at top of visualization. **(d)** Phylogenetic analysis of ∼15 kbp from at minimum two haplotypes per ape lineage and subsequent protein predictions for *MUC5B* haplotypes based on human exon boundary liftover. (*) = central node distinguishing species branches with 100 bootstrap support. Diagrams represent protein domains with the large central exon. **(e)** Scatterplot of total *MUC5B* exon 31 length (in base pairs) and total VNTR motif count across all VNTR domains in human and nonhuman primates. **(f)** Tiled alignments between representative haplotypes of each ape species (most common or most structurally unique haplotype per species) for *MUC5B. MUC5B* intron/exon boundaries distinguished by gene model at top of visualization.

Despite being more conserved in humans, there is extensive length variation among the protein coding MUC5B variants among great apes; however, cys domain copy number preceding the first tandem repeat is conserved. (Fig. 3d). Only orangutan and gibbon haplotypes harbor an additional cys domain that is distinctive from the other three cys domain types. We classify this sequence as a type V domain. Once again, orangutans carry the largest *MUC5B* VNTR domains (84-89 copies of the 29-mer). Excluding one haplotype from the Siamang gibbon, human alleles of *MUC5B* harbor shorter central exons on average with fewer VNTR total motifs compared to the NHP haplotypes (Fig. 3e) and little structural variation outside of the central exon (Fig. 3f).

### *MUC5AC* LD block structure and potential positive selection in East Asian populations

Given the important critical role of these mucins in the lung and gastric mucosa^1^ and the challenge associated with genotyping large VNTRs, we first investigated LD patterns among different human continental groups using D’. As expected, African populations showed the shortest LD blocks. Among non-African populations, a predominant single LD block corresponded to most of the MUC5AC protein coding gene (Fig. 4a). Because extended LD haplotypes are one potential signature of positive selection, we tested by simulation (Fig. 4b) if LD block sizes were significantly larger than the genome-wide distributions. When compared to population-specific distributions of LD block sizes in the 1KG dataset^19^, blocks in the *MUC5AC* region are large (top 5% distribution) in East Asians (n = 585) and Americans (n = 490) relative to Africans (n = 893), Europeans (n = 633), and South Asians (n = 601, Fig. 4b).

**Figure 4.**
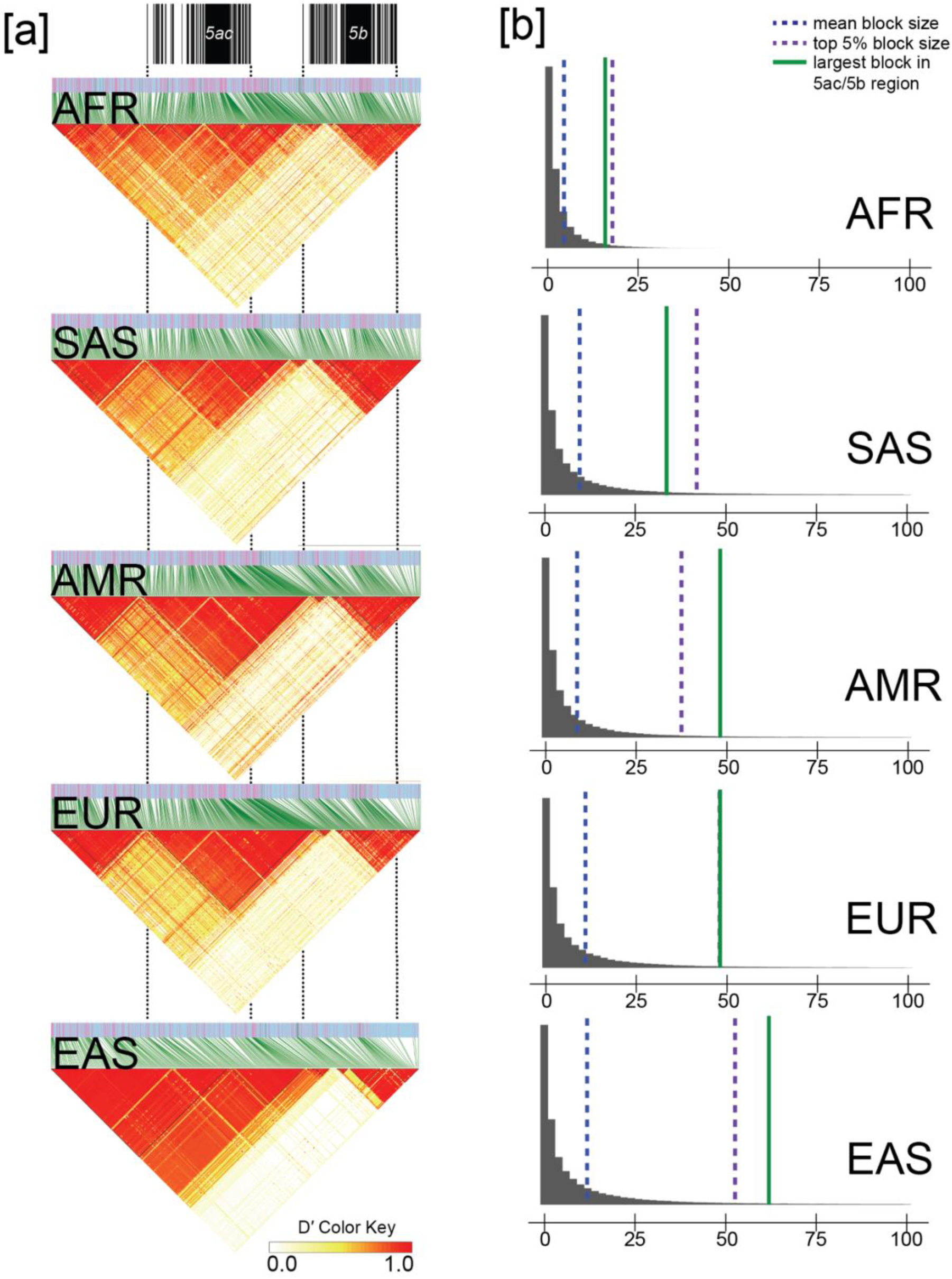
Linkage disequilibrium (LD) analysis of the *MUC5AC/5B* locus for African, American, European, East Asian, and South Asian genomes from the phased, short-read 1000 Genomes cohort. **(a)** LD plots for the *MUC5AC/5B* locus based on D’, with increasing red intensity indicative of higher LD between single-nucleotide polymorphisms (SNPs). AFR = African, SAS = South Asian, AMR = American, EUR = European, EAS = East Asian. **(b)** Autosome-wide LD block size distributions for each major population. Blocks above 100 kbp visually excluded as outliers (included in distribution analysis per population).

To test for positive selection more formally, we calculated Tajima’s D^37^ for 10 kbp segments spanning across *MUC5AC* and *MUC5B* in the 1KG sample set. We find a significant excess of rare variants for *MUC5B*-associated blocks in all super populations except Europeans, consistent with the action of positive selection (Table 1). Repeating the analysis for *MUC5AC*, only one population group (East Asians, Table 2) shows a negative Tajima’s D with the highest value corresponding to the 10 kbp segment immediately preceding the protein coding VNTR. East Asians are the only population with both an excess of rare variants and an abnormally large block of LD, thereby providing more compelling evidence of positive selection.

**Table 1.** Tajima’s D statistic for *MUC5B* in the 1KG.

| Pop | Gene | GRCh38 Chromosome 11 Bin |  |  |  |  |  |
| --- | --- | --- | --- | --- | --- | --- | --- |
|  |  | 1220000 | 1230000 | 1240000<sup>#</sup> | 1250000<sup>#</sup> | 1260000 | 1270000 |
| AFR | *MUC5B* | -1.09<br>(180) | -1.47<br>(162) | -1.84***<br>(444) | -1.87**<br>(279) | -1.94**<br>(209) | -2.10**<br>(248) |
| EUR | *MUC5B* | -0.44<br>(122) | -0.42<br>(81) | -1.49<br>(356) | -1.39<br>(191) | -1.56<br>(73) | -1.60<br>(103) |
| SAS | *MUC5B* | -0.55<br>(140) | -0.96<br>(100) | -1.76*<br>(328) | -1.77*<br>(188) | -1.59<br>(85) | -1.65<br>(102) |
| EAS | *MUC5B* | -0.25<br>(109) | -0.49<br>(62) | -1.58<br>(189) | -1.69*<br>(109) | -2.13**<br>(79) | -1.91*<br>(83) |
| AMR | *MUC5B* | -0.46<br>(130) | -1.22<br>(110) | -1.84<br>(353) | -1.87*<br>(201) | -1.95*<br>(108) | -1.90*<br>(122) |

Bin sizes of 10 kbp were used to compare values to the autosome-wide distribution per population in the 1000 Genomes Project<sup>19</sup> (1KG) cohort. (\*) Bottom 10% of autosome-wide Tajima's D values; (\*\*) Bottom 5% of autosome-wide Tajima's D values. Values in parentheses below each Tajima's D value correspond to the number of SNPs that were included in the calculation. (#) corresponds to bin containing VNTR sequence.

**Table 2.**
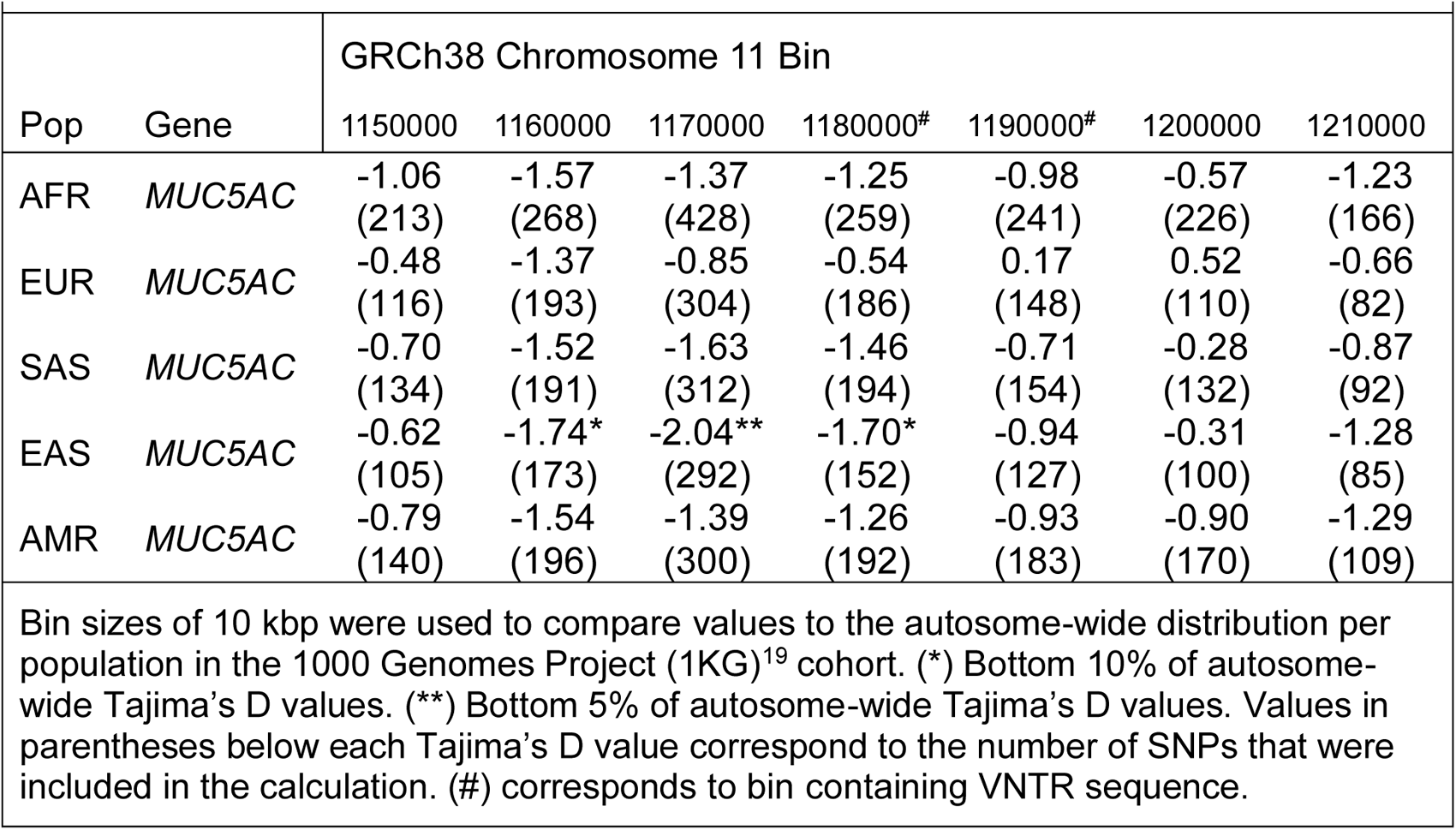
Tajima’s D statistic for *MUC5AC* in the 1KG.

| Table 2. Tajima's D statistic for *MUC5AC* in the 1KG |  |  |  |  |  |  |  |  |
| --- | --- | --- | --- | --- | --- | --- | --- | --- |
| Pop | Gene | GRCh38 Chromosome 11 Bin |  |  |  |  |  |  |
|  |  | 1150000 | 1160000 | 1170000 | 1180000<sup>#</sup> | 1190000<sup>#</sup> | 1200000 | 1210000 |
| AFR | *MUC5AC* | -1.06<br>(213) | -1.57<br>(268) | -1.37<br>(428) | -1.25<br>(259) | -0.98<br>(241) | -0.57<br>(226) | -1.23<br>(166) |
| EUR | *MUC5AC* | -0.48<br>(116) | -1.37<br>(193) | -0.85<br>(304) | -0.54<br>(186) | 0.17<br>(148) | 0.52<br>(110) | -0.66<br>(82) |
| SAS | *MUC5AC* | -0.70<br>(134) | -1.52<br>(191) | -1.63<br>(312) | -1.46<br>(194) | -0.71<br>(154) | -0.28<br>(132) | -0.87<br>(92) |
| EAS | *MUC5AC* | -0.62<br>(105) | -1.74*<br>(173) | -2.04**<br>(292) | -1.70*<br>(152) | -0.94<br>(127) | -0.31<br>(100) | -1.28<br>(85) |
| AMR | *MUC5AC* | -0.79<br>(140) | -1.54<br>(196) | -1.39<br>(300) | -1.26<br>(192) | -0.93<br>(183) | -0.90<br>(170) | -1.29<br>(109) |
| Bin sizes of 10 kbp were used to compare values to the autosome-wide distribution per population in the 1000 Genomes Project (1KG)<sup>19</sup> cohort. (*) Bottom 10% of autosome-wide Tajima's D values. (**) Bottom 5% of autosome-wide Tajima's D values. Values in parentheses below each Tajima's D value correspond to the number of SNPs that were included in the calculation. (#) corresponds to bin containing VNTR sequence. |  |  |  |  |  |  |  |  |

### tSNP discovery and short-read genotyping using Locityper

Given our sequence-resolved gene models and haplotype LD structure, we searched for tSNPs in high LD with VNTR haplogroups for the imputation of structural variants in short-read WGS datasets. To discover tSNPs, we encoded H1-H3 haplogroups as biallelic variants and tested for correlation (r^2^) with all SNPs within 10 kbp of the start and stop sites for *MUC5AC* (VNTR exon SNPs excluded). At a threshold of r^2^ > 0.85, we discovered 35 tSNPs for H1 (max r^2^ = 0.92), 5 tSNPs for H2 (max r^2^ = 0.89), and 52 tSNPs for H3 (max r^2^ = 1, Table S2). tSNPs for H1 are in moderate LD with H2 haplotypes (average r^2^ = 0.55) and those for H2 are in moderate LD with H1 haplotypes (average r^2^ = 0.64); however, tSNPs for H3 are in low LD with H1/H2 and make excellent imputation candidates for this group of variants (average H1/H2 r^2^ = 0.10). We found one tSNP distinguishing H1 and H2 of *MUC5B* that met our stringent r^2^ criteria (in GRCh38, chr11:1244757; H1 r^2^ = 0.0026 vs. H2 r^2^ = 1).

Next, we applied Locityper—a tool designed to genotype complex, multi-allelic loci like the VNTR-containing exons of *MUC5AC/5B*—to WGS datasets. Given a collection of high-quality reference alleles, such as those generated from long-read phased genomes, Locityper predicts the best pair of alleles for an unknown sample by examining read alignments and read-depth profiles across all allele pairs. Locityper has a short runtime allowing thousands of genomes to be rapidly characterized (https://github.com/tprodanov/locityper). We first tested the accuracy of Locityper in predicting haplogroup identities for *MUC5AC* and *MUC5B* in the HPRC/HGSVC genomes by performing LoO experiments. For *MUC5AC,* we estimated a genotyping accuracy of 95% for full diploid genotyping (both haplogroups correct) and 97.5% concordance for partial genotyping (one haplogroup correct), as compared to 97% and 98.5% for the closest available genotype (Methods, Fig. 5a, Table S4). For *MUC5B,* genotyping showed 100% accuracy in predicting the correct haplotype based on LoO experiments. (Fig. S4 and Table S3). Predictably, Locityper was less accurate in predicting protein isoforms due to homoplasy. For example, 91% and 81% of samples were correctly assigned to protein subgroups for *MUC5AC* and *MUC5B*, respectively (Table S3). It is likely that a larger sampling of reference haplotypes will significantly improve the future imputation of protein structure.

**Figure 5.**
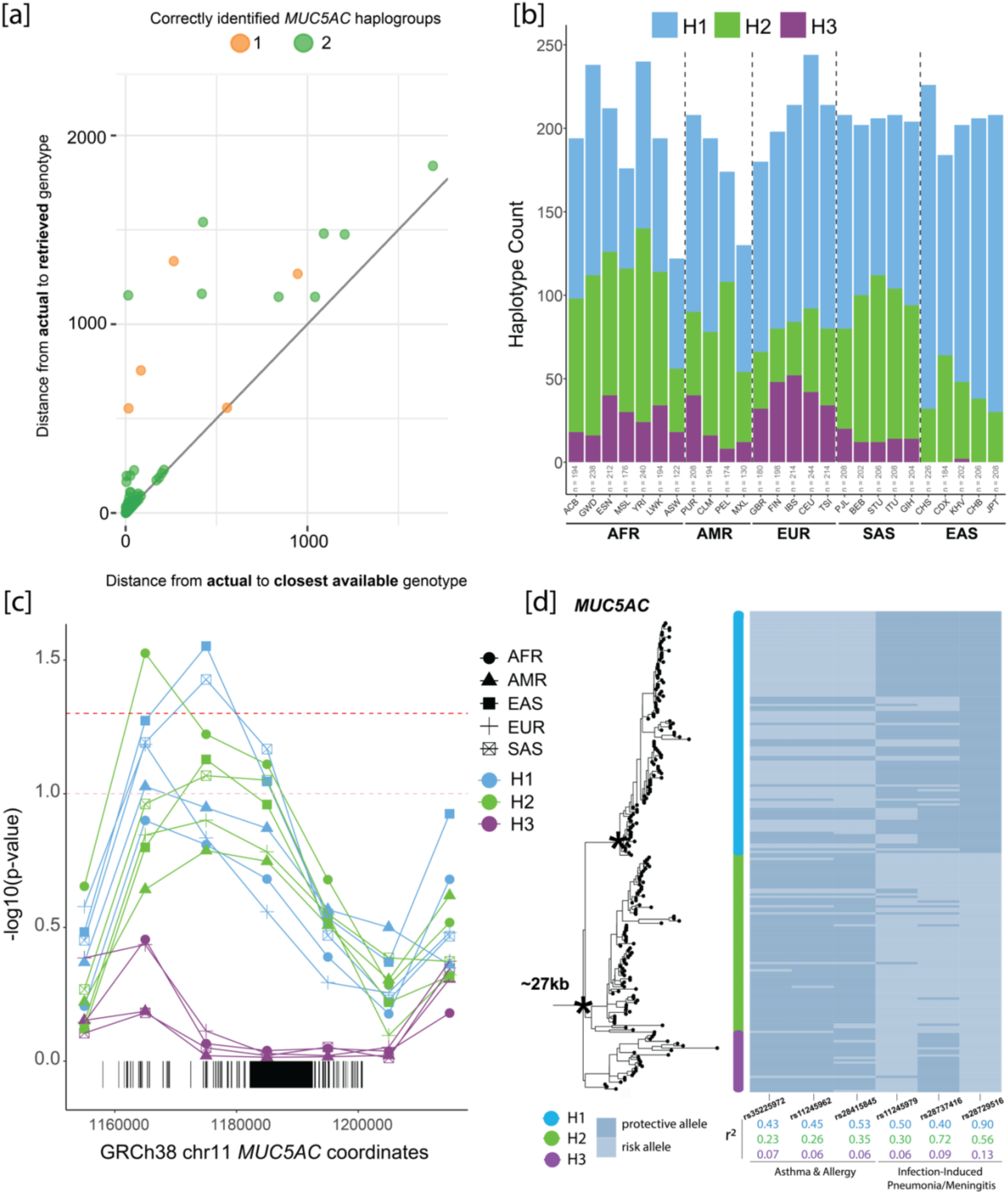
Genotyping of *MUC5AC* haplogroups with Locityper for population distributions and signatures of positive selection. **(a)** Locityper leave-one-out results comparing edit distances between actual vs. retrieved genotype (predicted from genotyper) versus edit distances between actual and closest possible genotype (best possible reference genotype from a multiple sequence alignment with true genotype) for *MUC5AC*. Dot color based on the number of haplotypes in diploid sample sets that were correctly genotyped. **(b)** *MUC5AC* haplogroup frequencies across superpopulations and populations in the 1KG dataset from Locityper predictions. AFR = African, AMR = American, EUR = European, SAS = South Asian, EAS = East Asian. ACB = African Caribbean in Barbados, GWD = Gambian in Western Division, ESN = Esan in Nigeria, MSL = Mende in Sierra Leone, YRI = Yoruba in Nigeria, LWK = Luhya in Kenya, ASW = Americans of African Ancestry in SW USA, PUR = Puerto Rican in Puerto Rico, CLM = Colombian in Colombia, PEL = Peruvian in Peru, MXL = Mexican Ancestry in Los Angeles USA, GBR = British in England and Scotland, FIN = Finnish in Finland, IBS = Iberian in Spain, CEU = Utah residents (CEPH) with Northern/Western European ancestry, TSI = Toscani in Italy, PJL = Punjabi in Pakistan, BEB = Bengali in Bangladesh, STU = Sri Lankan in the UK, ITU = Indian Telugu in the UK, GIH = Gujarati from Houston TX USA, CHS = Southern Han Chinese, CDX = Chinese Dai in China, KHV = Vietnamese in Vietnam, CHB = Han Chinese in Beijing, China, JPT = Japanese in Japan. **(c)** Significance of negative Tajima’s D values across the *MUC5AC* locus for genotyped haplogroups in each of the 1KG super populations relative to genome-wide distributions. The pink line corresponds to p-value of 0.1 and red line corresponds to p-value of 0.05. **(d)** Six genome-wide association (GWAS) risk and protective alleles mapped to the *MUC5AC* phylogeny. SNPs grouped based on disease association and squared correlations color-coded based on haplogroup partitioning.

Next, we genotyped all 2,600 unrelated genomes from the 1KG where a deeper short-read Illumina WGS dataset was recently generated. We compared haplogroup classification concordances for two high-confidence tSNPs with Locityper predictions for *MUC5AC* (H1 vs. H2/H3 tSNP: rs28542750, H3 vs. H1/H2 tSNP: rs769768817, Table S2). We found high concordance between the two methods, with 91% (n = 2,359) of the genomes yielding complete concordance for both haplotypes. For the remaining ∼9% (n = 241/2,600), most were discordant for only one of the two haplotypes (93%, n = 222/241) and differed for classification of H1 vs. H2 alleles (75%, n = 166/222). This is most likely because the highest confidence H1 tSNP is not as specific as that of the H3 tSNP used.

We leveraged the Locityper set of haplogroup predictions to consider population-specific representation of *MUC5AC* variants. In this larger population survey, we find that H2 is enriched in African genomes (47% of all African haplotypes) while H3 is found predominantly among African and European populations (18% and 21%, respectively; Fig. 5b). In sharp contrast, H3 is virtually absent among East Asians (0.37%); we identify only four haplotypes found exclusively among Vietnamese. It is interesting that among South Asians, H3 once again rises to common allele frequency (∼5%).

Using Locityper genotypes, we tested again for signatures of positive selection using Tajima’s D (Fig. 5c, Table 3). Our results suggest positive selection for both H1 and H2 *MUC5AC* haplogroups in East Asians and South Asians. We find that H2 in Africans yields more significantly negative Tajima’s D values in the regions of *MUC5AC* preceding the tandem repeat-containing exon, unlike the other super populations examined. In contrast, we find significant signatures of balancing selection (significantly positive Tajima’s D values) for *MUC5AC* H3 in Africans, Europeans, Americans, and South Asians. We tested for departure from Hardy-Weinberg equilibrium and found a significant depletion of homozygotes among Africans and Europeans (chi-squared test, AFR: p = 0.0368, EUR: p = 0.030), consistent with the action of balancing selection. These combined selection signatures suggest that while there is a likely immunological advantage to shorter haplotypes of *MUC5AC,* there has been a potential heterozygote advantage for the longer alleles (H3).

**Table 3.** Tajima’s D Statistic for *MUC5AC* stratified by Locityper haplogroups in the 1KG.

|  |  |  |  |  |  |  |  |  |  |
| --- | --- | --- | --- | --- | --- | --- | --- | --- | --- |
| Pop | Gene | Haplogroup | GRCh38 Chromosome 11 Bin |  |  |  |  |  |  |
|  |  |  | 1150000 | 1160000 | 1170000 | 1180000# | 1190000# | 1200000 | 1210000 |
| AFR | *MUC5AC* | H1 | -0.54<br>(156) | -1.36<br>(215) | -1.29<br>(321) | -1.18<br>(205) | -0.87<br>(197) | -0.47<br>(185) | -1.18<br>(131) |
|  |  | H2 | -1.17<br>(176) | -1.75**<br>(214) | -1.58*<br>(332) | -1.51*<br>(214) | -1.19<br>(199) | -0.71<br>(198) | -1.03<br>(142) |
|  |  | H3 | -0.19<br>(134) | -0.80<br>(166) | 0.19<br>(261) | 0.42*<br>(157) | 0.34*<br>(160) | 0.45*<br>(157) | -0.30<br>(104) |
| AMR | *MUC5AC* | H1 | -0.86<br>(121) | -1.67*<br>(155) | -1.60<br>(244) | -1.53<br>(165) | -1.18<br>(155) | -1.08<br>(141) | -0.84<br>(78) |
|  |  | H2 | -0.21<br>(85) | -1.09<br>(109) | -1.29<br>(193) | -1.24<br>(116) | -0.88<br>(117) | -0.45<br>(106) | -1.06<br>(78) |
|  |  | H3 | 0.42<br>(64) | 0.28<br>(70) | 1.61**<br>(142) | 1.79**<br>(94) | 1.72**<br>(93) | 1.58**<br>(69) | -0.10<br>(50) |
| EAS | *MUC5AC* | H1 | -0.67<br>(84) | -1.78*<br>(138) | -1.99**<br>(216) | -1.55*<br>(102) | -0.78<br>(84) | -0.41<br>(77) | -1.41<br>(77) |
|  |  | H2 | 0.81<br>(64) | -1.08<br>(112) | -1.53*<br>(198) | -1.32<br>(107) | -0.54<br>(99) | 0.32<br>(77) | -0.02<br>(54) |
|  |  | H3 | NA | NA | NA | NA | NA | NA | NA |
| EUR | *MUC5AC* | H1 | -0.84<br>(97) | -1.62*<br>(157) | -1.23<br>(257) | -0.81<br>(150) | -0.22<br>(125) | -0.11<br>(101) | -0.66<br>(77) |
|  |  | H2 | 0.83<br>(69) | -0.93<br>(105) | -1.02<br>(157) | -0.83<br>(109) | -0.40<br>(104) | 1.13<br>(73) | 0.16<br>(54) |
|  |  | H3 | -0.16<br>(75) | -0.28<br>(90) | 0.86<br>(188) | 1.75*<br>(110) | 1.93**<br>(103) | 1.38<br>(80) | -0.13<br>(51) |
| SAS | *MUC5AC* | H1 | -0.71<br>(108) | -1.66*<br>(159) | -1.85**<br>(257) | -1.64*<br>(153) | -0.74<br>(123) | -0.17<br>(111) | -0.74<br>(82) |
|  |  | H2 | -0.15<br>(97) | -1.36<br>(142) | -1.48*<br>(220) | -1.46*<br>(140) | -0.78<br>(113) | -0.46<br>(109) | -0.42<br>(72) |
|  |  | H3 | 1.20<br>(47) | 0.81<br>(58) | 1.65<br>(135) | 2.13**<br>(82) | 1.62<br>(90) | 2.39**<br>(54) | 0.27<br>(35) |
| Bin sizes of 10 kbp were used to compare values to the autosome-wide distribution per population and haplogroup in the 1000 Genomes Project<sup>19</sup> (1KG) cohort. (*) Bottom or top 10% of autosome-wide Tajima's D values. (**) Bottom or top 5% of autosome-wide Tajima's D values. Values in parentheses below each Tajima's D value correspond to the number of SNPs that were included in the calculation. (#) corresponds to bin containing VNTR sequence. Box shading indicate abnormally negative (light grey = bottom 10%, dark grey = bottom 5%) Tajima's D values relative to autosome-wide distributions within populations and haplogroups. Box outlines indicate abnormally positive (dashed box = top 10%, solid box = top 5%) Tajima's D values relative to the autosome-wide distribution within populations and haplogroups. |  |  |  |  |  |  |  |  |  |

### *MUC5AC* haplogroups in LD with GWAS risk SNPs and expression quantitative trait loci (eQTLs)

Since the tSNPs we discovered and characterized are unlikely to be directly genotyped in previous GWAS, we assessed the LD of *MUC5AC* haplogroups with risk and protective alleles that have been implicated in asthma/allergy phenotypes and infection-induced pneumonia/meningitis. The risk allele for three SNPs associated with the asthma/allergy phenotype (rs35225972^38^, rs11245962^39^, and rs28415845^40^; European cohorts) are in moderate LD with H1 variants of *MUC5AC* (Fig. 5d). Conversely, the protective alleles for two SNPs associated with infection-induced pneumonia/meningitis (rs11245979^41^, rs28729516^43^; European cohorts) are in higher LD with H1, with the protective allele for rs28729516 acting as a tSNP for this haplogroup. Surprisingly, a GWAS-risk SNP for severe tuberculosis-induced meningitis was in moderately high LD with H2 variants (Vietnamese cohort, rs28737416^42^), i.e., the haplogroup showing a modest signature of positive selection among East Asians. Similarly, we examined SNP-associated eQTLs for *MUC5AC* identified in the upper airways of African American and Hispanic children^55^. These eQTLs were parsed into two independent groups related to either increased (group A) or decreased (group B) *MUC5AC* expression. We found that group A eQTLs (increased *MUC5AC* expression/decreased lung function) have an average r^2^ of 0.79 for the risk variant and H1 *MUC5AC* alleles, whereas group B eQTLs (decreased *MUC5AC* expression/asthma protective) correlate (r^2^=0.82) with the H3 *MUC5AC* alleles (Table S4). Combined, these findings suggest that differences in VNTR structure are likely important considerations in both disease susceptibility as well as mucin expression in asthma.

### MUC5AC and MUC5B PheWAS in All of Us

To identify phenotypes associated with secreted airway mucin genetic variation, we performed a PheWAS using data from *All of Us*^47^ (n = ∼165,150). We first validated our analytical pipeline by testing for a known disease association with the *MUC5B* regulatory polymorphism and interstitial lung diseases^48^. We find significant associations after Bonferroni correction for rs35705950 in all samples (including age, sex, and PCs1-3 as covariates) for respiratory phenotypes, including International Classification of Diseases (ICD) codes for alveolar and parietoalveolar pneumonopathy (p = 6.89E-44, OR = 2.05), idiopathic fibrosing alveolitis (p = 2.14E-36, OR = 2.85), postinflammatory pulmonary fibrosis (p = 2.62E-34, OR = 1.82), extrinsic allergic alveolitis (p = 3.96E-08, OR = 2.38), bronchiectasis (p = 9.77E-7, OR = 1.32), and pulmonary congestion and hypostasis (p = 2.25E-05, OR = 1.30; Fig. S5). When we assessed phenotype associations within individual populations, we found significant associations with two or more of the same respiratory phenotypes in admixed Americans and Europeans (Table S5). We did not assess disease associations with the *MUC5B* tSNP distinguishing H1 and H2 because the predominant protein variants in these haplogroups differ by only one VNTR codon (unlikely to have phenotypic effects).

We find no correlated phenotypes that survive multiple hypothesis testing correction for *MUC5AC* H1, H2, and H3 tSNPs (Methods). It is interesting to note, however, that H3 tSNPs approached significance for protection against degeneration of the macula and the posterior pole of the retina (p = 1.76E-4 – 9.49E-4, OR=0.91; Table S6). We repeated the analysis separately for heterozygotes and homozygotes at rs36151150 (*MUC5AC* H3 tSNP) and find increased significance for the protective phenotype among heterozygotes, despite a reduction in alleles upon removal of homozygotes (heterozygous: p = 2.41E-4, OR = 0.89; homozygous: p = 0.145, OR = 0.94; Table S6).

## DISCUSSION

Using numerous high-quality long-read genome assemblies, we performed the first population-level genetic diversity survey of *MUC5AC* and *MUC5B* protein structural polymorphism. The long, highly variable, protein coding VNTRs of both loci precluded the study of these genes from short-read WGS datasets. Initial efforts to resolve these loci using long-read sequencing have relied on platforms with higher error rates (PacBio Continuous Long Reads) and have been limited to only a few individuals (n=4)^7^; however, recent advances in long-read sequencing technologies and *de novo* genome assembly algorithms^11,15,16^ have made complete characterization of these genes possible for the first time^9,10^. Characterization of these structurally diverse loci opens a path to improved understanding of this form of human genetic diversity in health and disease.

While our results recapitulate the long-held belief that the *MUC5B* VNTR is less variable than other secreted mucins^51^, we have identified new protein-encoding structural variants of likely functional consequence among Africans, demonstrating that this protein is not intolerant to variation. This is perhaps not surprising given the greater genetic diversity expected among Africans^56^. These variants have likely been missed because most studies of *MUC5B* polymorphism have been conducted within European populations (e.g., *MUC5B* promoter polymorphism^48^). This demonstrates the importance of initiatives from consortia like the HGSVC^9^, HPRC^10^, and *All of Us*^47^ to broadly survey human genetic diversity with high-accuracy, long-read sequencing platforms.

In contrast to *MUC5B*, we discovered extensive amino-acid composition and size variation within the large central exon of *MUC5AC.* This difference may be related to their varying functional roles. *MUC5B,* for example, is constitutively expressed throughout the airways, while *MUC5AC* is predominantly expressed in the upper airways and is highly responsive to inflammation^57^. Therefore, *MUC5AC* has likely evolved independently from *MUC5B* to respond to a wider variety of pathogenic challenges^1^. Much of the protein structural polymorphism in MUC5AC is associated with cys domain and tandem repeat domain copy number. The consistent presence of cys domains separating distinct tandem repeat domains in both *MUC5AC* and *MUC5B* for humans and NHPs could function as a prevention method for excessive VNTR expansion at these loci.

Our comparative analyses with NHPs also indicates that VNTR length has generally decreased in the human lineage over the course of ape evolution for both genes. Increased VNTR length and subsequent glycosylation is predicted to enhance the interaction of the mucins with water^56^ (thereby altering the mucus biophysical properties) and an increased number of cys domains may enhance non-covalent self-interactions that make the gel impermeable^59^. Thus, it is possible that longer variants of both mucins contribute to changes in the viscoelastic properties of mucus that contribute to disease phenotypes, such as asthma and cystic fibrosis. In this regard, it is noteworthy that respiratory disease is a particularly pervasive problem affecting NHPs in captivity^60^; therefore, the reduction in overall VNTR length (especially in H1 and H2 haplogroups) may have been particularly adaptive in humans. Because of our detailed curation of many *MUC5AC* and *MUC5B* human haplotypes^9,10,19^, further experimental work uncovering how *MUC5AC* and *MUC5B* protein length variation could impart functional differences for airway disease and pathogen entrapment is now possible.

Within the human population, we distinguish three major *MUC5AC* haplogroups (H1-H3) that generally correlate with VNTR length (H3 encoding the longest and H1 the shortest MUC5AC molecules, Fig. 1). The longer haplogroup variants are significantly depleted among genomes of East Asian descent. We observe signatures of positive selection for the two most common (and shorter) haplogroups, H1/H2, as evidenced by an excess of rare variants (Tajima’s D) and extended LD consistent with an immunological advantage for shorter *MUC5AC* alleles. We find that the strongest signatures of positive selection are among East Asian samples. While this could be in part due to recent population bottlenecks/rapid population expansion^61^, our genome-wide survey of LD blocks in East Asian genomes places *MUC5AC* block length in the top 5% (Fig. 4). These findings may be relevant to the increased prevalence of asthma in individuals of European and African American descent when compared to Asians^62,63^, although many other mitigating factors, such as environmental exposures^62^, play an important role.

We leveraged the LD and structural variation differences present within the 206 assembled haplotypes of MUC5AC and MUC5B to genotype short-read WGS data from the same 1KG samples. Using the recently developed program Locityper, we estimate a high degree of genotyping accuracy (∼95% based on LoO experiments) with most of the error associated with false predictions of the cys copy number preceding the first large tandem repeat domain. Applying Locityper to the high-coverage WGS data recently generated from 2,600 1KG samples^19^ confirms the striking population stratification and signature of positive selection among East Asian populations (Fig. 5). Given the importance of *MUC5AC* as a well-known genetic modifier of epithelial diseases like cystic fibrosis^8^ and asthma/allergy^38,39,40^, it will be critical to continue cataloging greater haplotype diversity and improving long-read to short-read genotyping assays at this locus.

Our study of the genetic diversity of MUC5AC suggests different forces of both balancing and positive selection are operating. Unlike H1 and H2, where LD block size and Tajima’s D suggest positive selection, our analysis of H3 provides preliminary evidence of heterozygote advantage based on positive Tajima’s D among Europeans and Africans. The molecular basis for this is unknown, but it is interesting that a protective effect was suggested by PheWAS for macular/retinal degeneration and enriched in H3 heterozygotes (Table S6). Given that *MUC5AC* expression has been previously associated with dry eye syndrome^64,65^, it is feasible that H3 variants provide a protective function against ocular disease. The mucin genes (including *MUC5AC* and *MUC5B*) are expressed in numerous epithelial tissues outside of the lungs; therefore, the signatures of selection we have uncovered may be due to more than just lung traits. It will be critical to understand these biological nuances and control for population substructure in future genome-wide associations involving much larger sample sizes.

At a broader level, the strategy we outlined will be applicable to other mucin loci and biomedically important gene families. There are numerous genes with protein-encoding structural polymorphisms that have generally been excluded from surveys of human genetic variation. Some of these are already known to contribute significantly to human disease, such as *LPA*^66^ and *CYP2D6*^67^, while others are suggestive, such as *HRNR*^68^. Accurate and contiguous sequence haplotypes of these polymorphic genes from long reads as part of the HGSVC, HPRC, and *All of Us* will facilitate complete protein domain sequence curation, LD block structure analysis, and tSNP identification. Importantly, the resulting panels of sequence-resolved haplotypes, coupled with structural genotyping tools such as Locityper, could facilitate direct genotyping from short reads in large population cohorts like *All of Us* or the UK Biobank. As long-read sequencing methods continue to be optimized and become less expensive in the coming years, the importance of these more complex forms of human genetic variation and their relevance to human disease will become realized.

## Supporting information

Supplemental_Figs_and_SmallTables

## ACKNOWLEDGMENTS

We thank Tonia Brown and Michelle D. Noyes for assistance in editing this manuscript. We thank the HGSVC for access to the underlying PacBio HiFi CCS reads for local assembly for the *MUC5AC/5B* locus and the primate T2T project, especially Kateryna Makova and Adam Phillippy, for access to the high-quality data ape genome assemblies via GenomeArk. We thank Brian Browning, Devin Schweppe, and Nick Riley for their intellectual contributions to the experimental designs and visualizations contained in this manuscript. This work was supported, in part, by US National Institutes of Health (NIH) grants HG002385, HG010169, and HG007497 to E.E.E. E.E.E. and J.D.B. are investigators of the Howard Hughes Medical Institute. This article is subject to HHMI’s Open Access to Publications policy. HHMI lab heads have previously granted a nonexclusive CC BY 4.0 license to the public and a sublicensable license to HHMI in their research articles. Pursuant to those licenses, the author-accepted manuscript of this article can be made freely available under a CC BY 4.0 license immediately upon publication.

## AUTHOR CONTRIBUTIONS

E.G.P, P.H., and E.E.E. conceived and planned the experiments. W.K.O., P.H., and J.D.B. provided critical intellectual support during project design. General methodologies were conceived by E.G.P., P.H., A.S., and E.E.E. E.G.P. performed data curation and formal analyses. T.P. and T.M. supported E.G.P. with Locityper analyses and data visualizations. E.N., E.J.K., and P.N.V. conceived and performed the PheWAS analyses. W.T.H. and K.M.M. provided technical and scientific consultation. Data visualization was designed by E.G.P and E.E.E. E.G.P. and E.E.E. wrote the original manuscript, with edits and reviews from all authors. All authors provided critical feedback that shaped the research and analysis outlined in this manuscript.

## DATA AND CODE AVAILABILITY

The assemblies generated for this project (not previously published by the HGSVC^9^) will be uploaded and accessioned via IGSR after initial submission.

