## Supplemental_Figs_and_SmallTables for "Structural and genetic diversity in the secreted mucins, *MUC5AC* and *MUC5B*"

**Note S1.** Using tandem repeats finder, we discovered an 8-mer VNTR with a canonical motif of TCACCCAC in all human haplotypes. Haplogroup 3 (H3) alleles harbor more copies of the repeat and a lower percent motif identity compared to haplogroup 1 and 2 alleles in humans. STREME (Sensitive, Thorough, Rapid, Enriched Motif Elicitation) analysis reveals five 24-mers that are exclusive to H3 alleles for genotyping this haplogroup (ACCATTCACTCACCCATTCACCCATTCACC, ACTCACCC ACTCACCCATTCACCCA TTCAC, ACTCACCCACTCACTCACTCACCTACTCAA, CAG TGGGTGATTGAGTG GGTGAATGGGTGA, and TAAGTTGAGTGAGTGAGGTGAGTG AGTGGA). The canonical 8-mer motif is exclusive to human haplotypes, with all NHPs featuring motif lengths of 12-28 nucleotides. Total VNTR copy number is also highly variable among the nonhuman apes, leading to differences in total sequence length for intron 15 across haplotypes (Fig. S2).

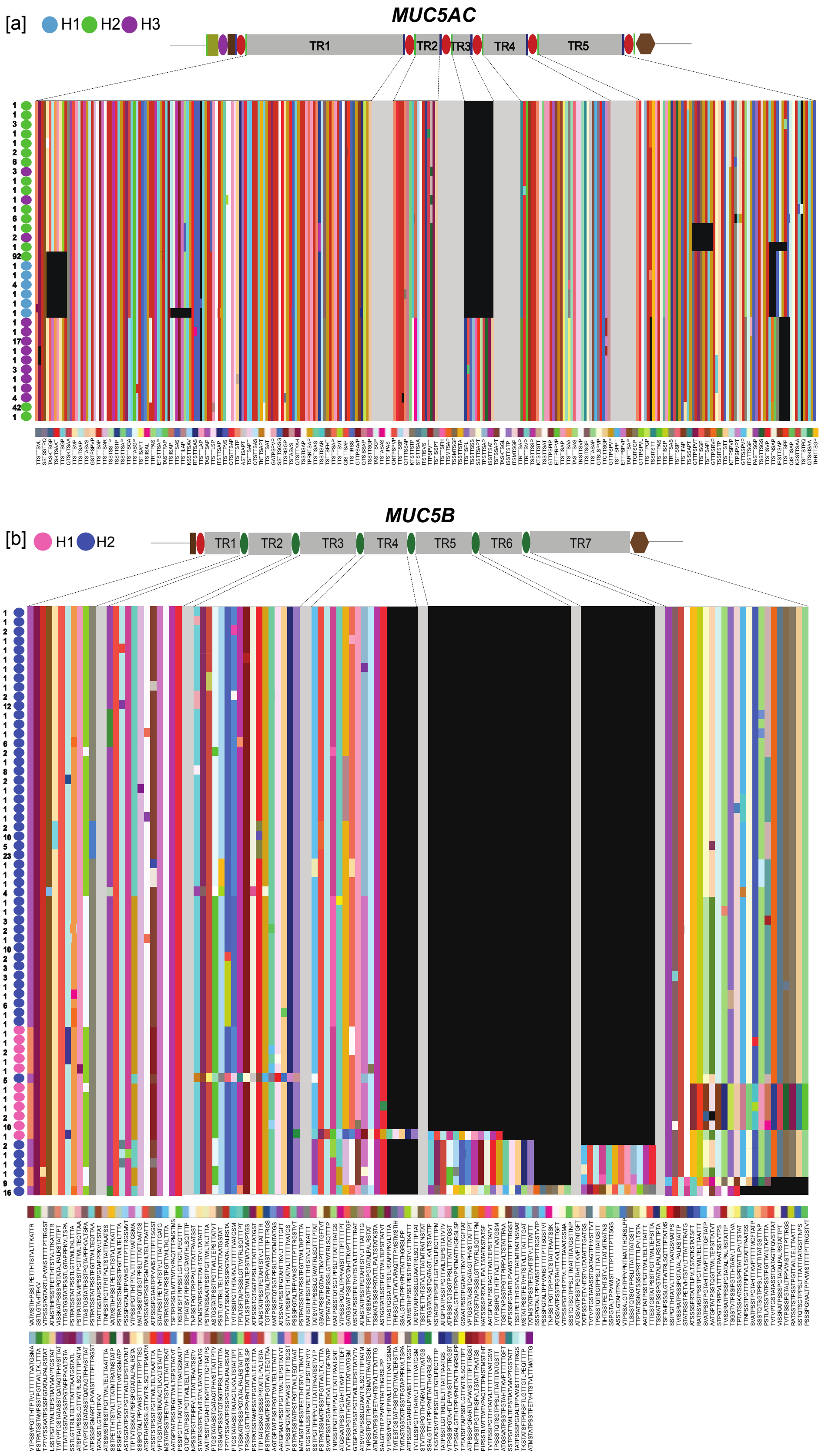
[a]
H1
H2
H3
MUC5AC
TR1
TR2
TR3
TR4
TR5
[b]
H1
H2
MUC5B
TR1
TR2
TR3
TR4
TR5
TR6
TR7

**Figure S1. MUC5AC and MUC5B protein VNTR motif utilization across 206 human haplotypes. (a)** MUC5AC protein 8-mer variation plots across the five possible VNTR domains. Domain plot height corresponds to the total number of unique alleles across the large central exon. Numbers per unique allele correspond to the number of haplotypes that are matching in motif composition across all domains in linear sequence space. Black boxes indicate an absence of sequence. Gray represents cys domain sequence between distinct VNTR domains. Alleles are ordered vertically by hierarchical clustering of motif composition. Motifs are shifted in sequence space based on the same identity (indicating linear shifts of individual motifs). **(b)** MUC5B protein 29-mer variation plots across the five possible VNTR domains. Domain plot height corresponds to the total number of unique alleles across the large central exon. The numbers per unique allele correspond to the number of haplotypes that are matching in motif composition across all domains in linear sequence space. Black boxes indicate an absence of sequence. Gray represents cys domain sequence between distinct VNTR domains. Alleles are ordered vertically by hierarchical clustering of motif composition. Motifs are shifted in sequence space based on the same identity (indicating linear shifts of individual motifs).

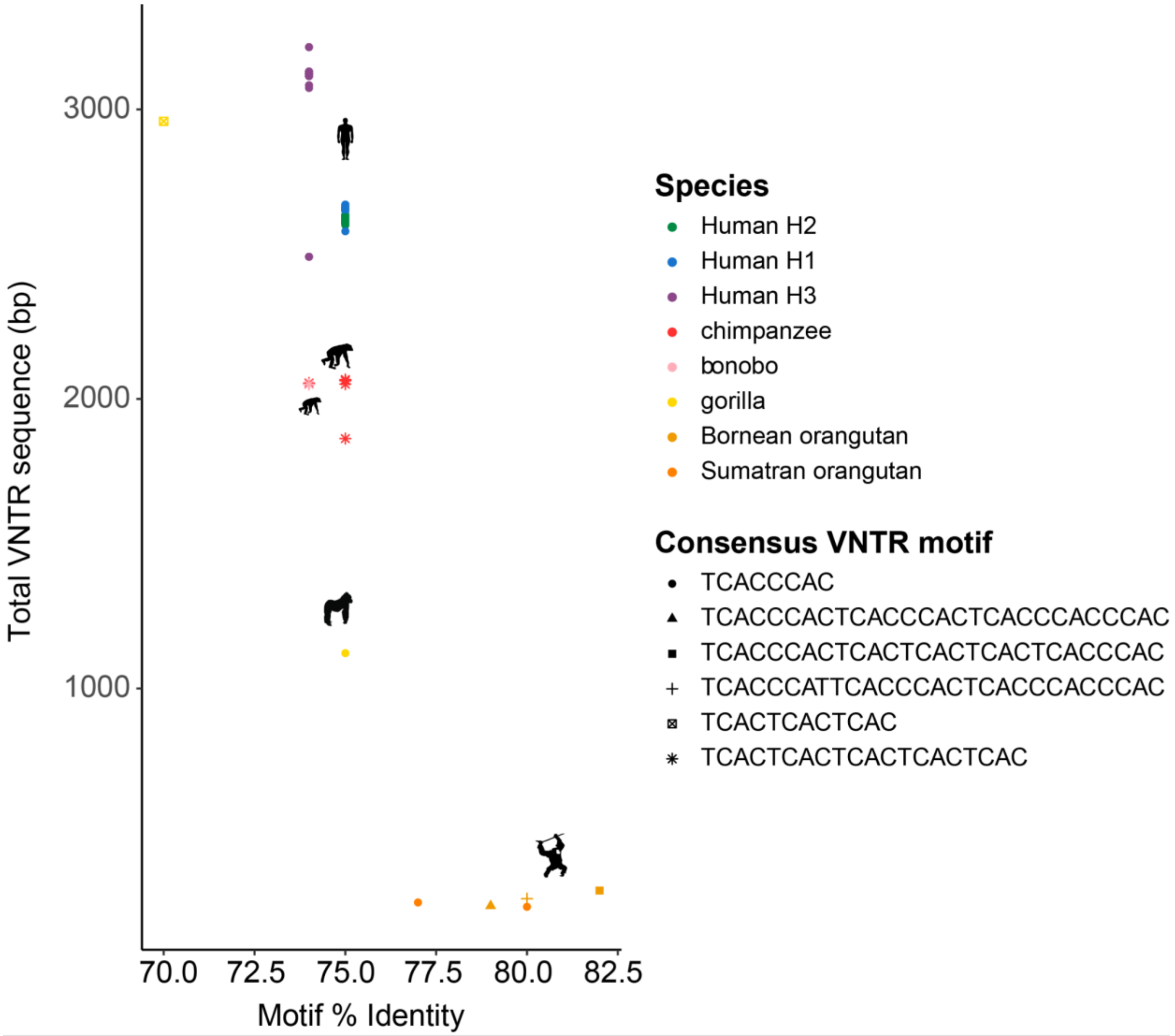


**Figure S2. Intron 15 VNTR sequence characterization in *MUC5AC* for humans and nonhuman apes.** Comparison of total VNTR sequence length (in base pairs) and canonical motif percent identity across 206 human haplotypes and at least two haplotypes per nonhuman ape species. Species/haplogroup identity coded by color and shape coded by motif.

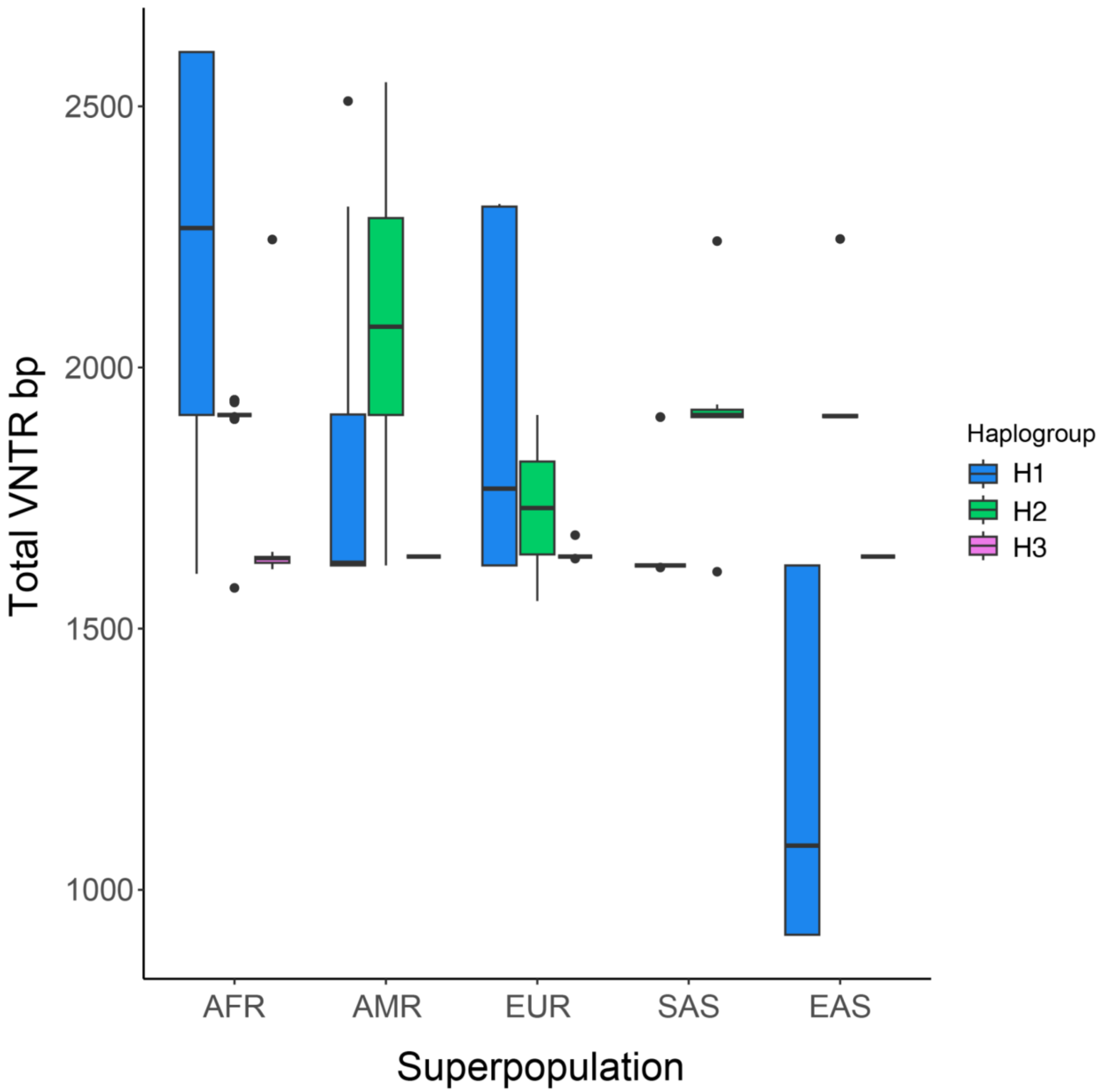


**Figure S3.** ***MUC5AC*** **VNTR enhancer polymorphism across human super populations in the HGSVC/HPRC sample set, stratified by haplogroup identity.** Total VNTR bp indicates the longest continuous track of degenerate 8-mer VNTR sequence in the enhancer region of *MUC5AC*. Haplogroups correspond to phylogenetic clades of the *MUC5AC* locus.

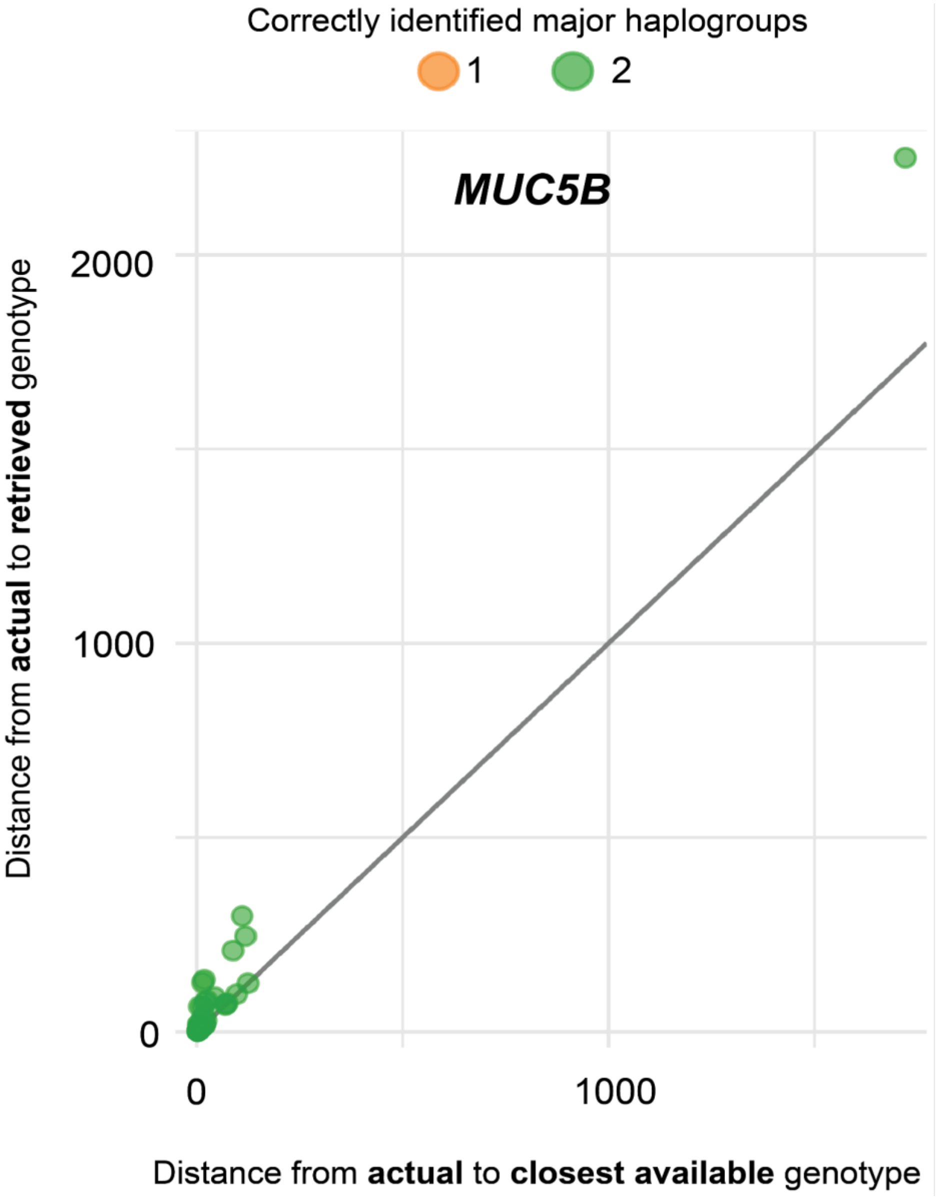


**Figure S4. Genotyping accuracy of *MUC5B* haplogroups with Locityper.** Locityper leave-one-out results comparing edit distances between actual and retrieved genotype (predicted from genotyper) versus edit distances between actual and closest possible genotype (best possible reference genotype from multiple sequence alignment with true genotype) for *MUC5B*. Dot color corresponds to the number of haplotypes in diploid sample sets that were correctly genotyped.

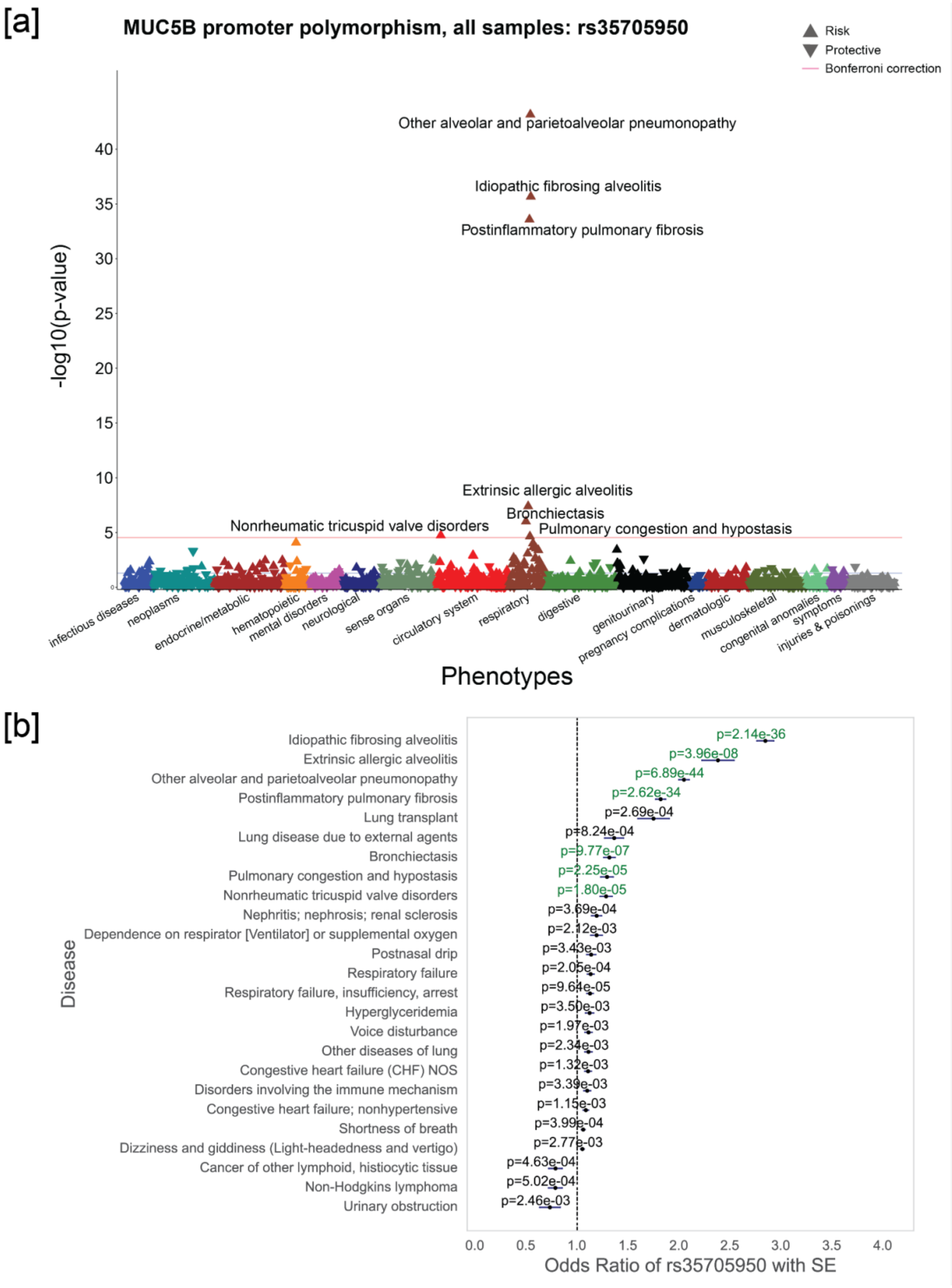
[a]
MUC5B promoter polymorphism, all samples: rs35705950
Risk
Protective
Bonferroni correction
-log10(p-value)
40
35
30
25
20
15
10
5
0
Other alveolar and parietoalveolar pneumonopathy
Idiopathic fibrosing alveolitis
Postinflammatory pulmonary fibrosis
Extrinsic allergic alveolitis
Bronchiectasis
Nonrheumatic tricuspid valve disorders
Pulmonary congestion and hypostasis
infectious diseases
neoplasms
endocrine/metabolic
hematopoietic
mental disorders
neurological
sense organs
circulatory system
respiratory
digestive
genitourinary
pregnancy complications
dermatologic
musculoskeletal
congenital anomalies
symptoms
injuries & poisonings
Phenotypes
[b]
Disease
Idiopathic fibrosing alveolitis
Extrinsic allergic alveolitis
Other alveolar and parietoalveolar pneumonopathy
Postinflammatory pulmonary fibrosis
Lung transplant
Lung disease due to external agents
Bronchiectasis
Pulmonary congestion and hypostasis
Nonrheumatic tricuspid valve disorders
Nephritis; nephrosis; renal sclerosis
Dependence on respirator [Ventilator] or supplemental oxygen
Postnasal drip
Respiratory failure
Respiratory failure, insufficiency, arrest
Hyperglyceridemia
Voice disturbance
Other diseases of lung
Congestive heart failure (CHF) NOS
Disorders involving the immune mechanism
Congestive heart failure; nonhypertensive
Shortness of breath
Dizziness and giddiness (Light-headedness and vertigo)
Cancer of other lymphoid, histiocytic tissue
Non-Hodgkins lymphoma
Urinary obstruction
p=2.14e-36
p=3.96e-08
p=6.89e-44
p=2.62e-34
p=2.69e-04
p=8.24e-04
p=9.77e-07
p=2.25e-05
p=1.80e-05
p=3.69e-04
p=2.12e-03
p=3.43e-03
p=2.05e-04
p=9.64e-05
p=3.50e-03
p=1.97e-03
p=2.34e-03
p=1.32e-03
p=3.39e-03
p=1.15e-03
p=3.99e-04
p=2.77e-03
p=4.63e-04
p=5.02e-04
p=2.46e-03
0.0
0.5
1.0
1.5
2.0
2.5
3.0
3.5
4.0
Odds Ratio of rs35705950 with SE

**Figure S5. PheWAS of the *MUC5B* promoter polymorphism rs35705950 in *All of Us*. (a)** Phenome-wide association study (PheWAS) Manhattan Plot of phenotype groupings and associations with the *MUC5B* promoter polymorphism rs35705950 in all populations (i.e., all samples). Only phenotypes that passed Bonferroni correction are noted by name. **(b)** Odds ratios of the top 25 diseases (based on ordered p-values) for associations with rs35705950. Green indicates phenotypes that passed Bonferroni correction. SE = standard error (horizontal bars corresponding to each p-value).

| Table S3. | Locityper leave-one-out haplogroup and protein group concordance results in for *MUC5AC* and *MUC5B* | | |
|---|---|---|---|
| Gene | Concordance Comparison | Haplogroup Concordance | Protein Group Concordance |
| *MUC5AC* | Full genotype match: retrieved vs. actual | 95% (94/99) | 91% (90/99) |
| | Full genotype match: closest available vs. actual | 97% (96/99) | 91% (90/99) |
| | Partial genotype match: retrieved vs. actual | 97.5% (193/198 | 95.5% (189/198) |
| | Partial genotype match: closest available vs. actual | 98.5% (195/198) | 95.5% (189/198) |
| *MUC5B* | Full genotype match: retrieved vs. actual | 100% (99/99) | 81% (80/99) |
| | Full genotype match: closest available vs. actual | 100% (99/99) | 95% (94/99) |
| | Partial genotype match: retrieved vs. actual | 100% (198/198) | 89% (176/198) |
| | Partial genotype match: closest available vs. actual | 100% (198/198) | 97.5% (193/198) |

Table S1. See excel file titled "Table_S1.xlsx"

Table S2. See excel file titled "Table_S2.xlsx"

Table S4. See excel file titled "Table_S4.xlsx"

Table S5. See excel file titled "Table_S5.xlsx"

Table S6. See excel file titled "Table_S6.xlsx"
